# Evolution of the motor cortex microstructure and its lateralization: a comparative study of chimpanzees and humans

**DOI:** 10.64898/2026.08.20.745984

**Authors:** M. Chauvel, E. Kirilina, I. Lipp, F. Buettner, C. Jäger, K.J. Pine, L.J. Edwards, S.J. Ebel, K.S. Kopp, S. Helbling, P. McColgan, D. Rose, T. Gräßle, EBC Consortium, R. McElreath, D. Chaimow, C. Crockford, R.M. Wittig, N. Weiskopf

## Abstract

Human hand coordination exceeds that of other species, including great apes, and is marked by pronounced right-hand dominance. This specialization parallels an expansion of its cortical representation, forming the hand-knob in the motor cortex. In humans, this region shows high myelination on quantitative MRI (qMRI), but whether this feature is shared with great apes remains unclear. It is also unknown whether increased right-hand dominance in humans is mirrored by greater hemispheric asymmetry in cortical microstructure. Using high-resolution qMRI, we compared motor cortex subdivisions controlling the leg, hand, and face in humans and chimpanzees. We found consistently higher myelin and iron content in the hand-knob in both species, suggesting an evolutionarily conserved role. However, only humans showed enhanced rightward lateralization. These results highlight both conserved and species-specific features of the motor cortex, offering insights into the evolution of manual dexterity and handedness.

## Introduction

The unparalleled manual dexterity of humans represents a major evolutionary leap, setting Homo sapiens apart from both our hominin ancestors and extant great apes^1^. Even for our close relatives, chimpanzees, who exhibit remarkable hand dexterity combining the functional requirement for both arboreal locomotion and tool-use manipulation, the ability to perform human-like fine hand movements remains limited^2–4^. Manual dexterity developed alongside *handedness*, a preference for one hand over the other, with approximately 90% of human individuals favoring their right hand for skilled tasks^5–7^. Interestingly, the hand preference in chimpanzees at the population level is consistently weaker compared to those observed in humans^8^ and remains the subject of ongoing debate. While some studies have reported a higher prevalence of right-hand preference at the population level in some tasks^9–11^, others suggest that hand preference in chimpanzees is individual or task dependent^12,13^. Human exceptional capacity for fine motor control and complex tool use, particularly with the right hand, is attributable not only to different hand anatomy but also to the sophisticated neural processes governing hand movements.

The primary motor cortex (M1, Brodmann Area 4), located anteriorly to the central sulcus in the frontal lobe, plays a key role in hand movement control. As part of a broader motor system that includes premotor areas and supplementary motor areas, M1 generates neural impulses that control the planning and execution of voluntary movements^14^ via the cortico-spinal tract (CST)^15^. M1 is not only functionally but also anatomically structured in a somatotopic manner, meaning that different regions (or cortical fields) of M1 are involved in voluntary movements of different parts of the body^16,17^.

Notably, the central sulcus in hominoids exhibit a “hand-knob”, an anatomical feature hosting the subpart of M1 controlling hand movements^16,18,19^. Located between the area controlling lower extremities^20^ and areas associated with oro-facial motor movements^21^ (see also Figure 1), the hand-knob is one of the few discernible gross anatomical landmarks in the human brain that shows a highly consistent functional role across individuals^22^. The hand-knob was also identified and studied in chimpanzees and other great apes^23,24^ but appears to be either absent or indistinct in monkeys^25^. Electrophysiological and lesion studies in the early to mid-1900s^26–28^ confirmed that, similar to humans, the hand-knob controls hand movement in chimpanzees^29,30^.

**Figure 1.**
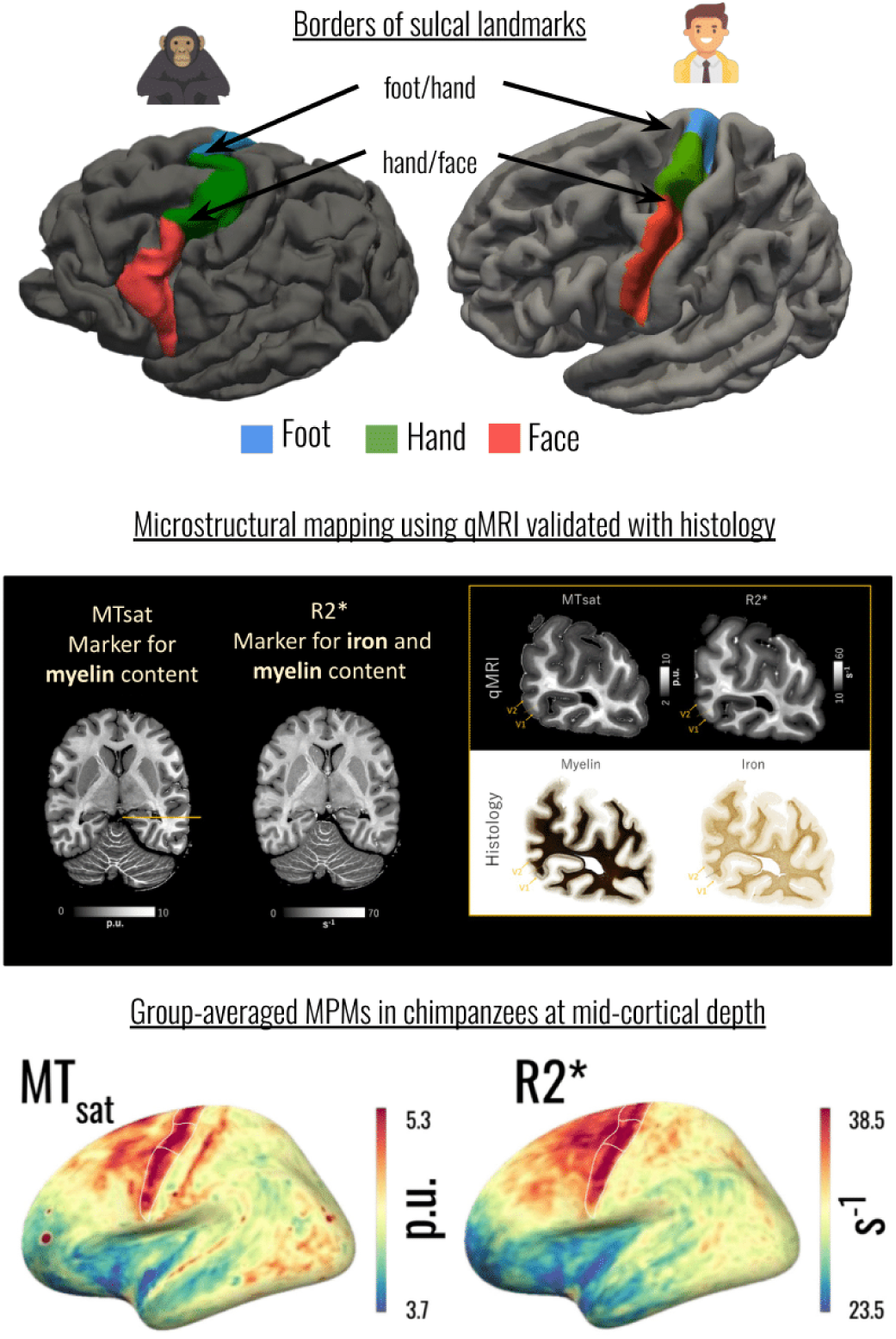
Motor sulco-gyral landmarks and quantitative MRI maps. Top row: labeling of the different motor cortical fields representing the different body parts along the central sulcus. Red, green and blue labels correspond respectively to face, hands and foot areas. Labels are superimposed on the chimpanzee and human averaged 3D pial meshes. Middle row: examples of MTsat and R2* quantitative MRI maps, highlighting differences in myelin and iron accumulation in the occipital cortex and the correspondence found between qMRI and histology (Gallyas stain for myelin and Perls’s stain for iron) (modified from Lipp et al.). Bottom row: Cortical surface with projected MTsat and R2* values for chimpanzees at mid-cortical depth with overlaid cortical fields delineation (white lines).

A remarkable feature of the hand-knob in humans is its exceptionally high levels of cortical myelin, compared to other subdivisions of M1^31^, despite sharing the same cytoarchitecture. It remains unclear if the exceptional myelination of hand knob in humans is a species-specific adaptation or evolutionary conserved feature present already in great apes. Moreover, the question arises, if handedness - a recently evolved feature of motor behaviour - is reflected by cortical lateralization during the evolution of the hominid lineage. Interestingly, findings on macroanatomical asymmetry of the human motor cortex and its relationship with handedness in humans remain inconsistent. Some studies reported asymmetries in the precentral gyrus and a deeper central sulcus in the hemisphere contralateral to the dominant hand^32^, while larger-scale analyses found no significant differences in cortical morphology between left- and right-handed individuals^33,34^. In chimpanzees, no population level asymmetry in the size of the hand-knob was observed, although some studies suggest associations between individual hand preference and sulcal morphology^35,36^.

At the histological level, results are similarly mixed. In humans, cytoarchitectonic studies by^37^ on the neuropil found some differences in Brodman area 4 (M1) implying higher neuropil density in the left motor cortex in humans, while other studies did not report significant interpretable differences^38^. In chimpanzees, only one study reported limited evidence in asymmetries in neuron density within specific cortical layers of the hand-knob^11^. Investigations of white matter bundles such as the cortico-spinal tract (CST) revealed no significant correlation between leftward-asymmetry and handedness in humans on the population-level^39,40^ or in chimpanzees^41^. No histological study systematically addressed the relationship between lateralization of cortical myelination and handedness in humans and chimpanzees. This may be attributed to the fact that myelin quantification using histology is highly challenging, particularly when comparisons must be made across macroscopic and spatially distant areas. Histological methods for myelin detection, which rely on stains, are limited by variability in staining efficiency, are not quantitative and are not readily applicable in three dimensions or across large populations.

Recent advances in ultra-high-resolution quantitative magnetic resonance imaging (qMRI) offer a unique opportunity to characterize important features of cortical microstructure non-invasively. Quantitative MRI parameters are sensitive to tissue microstructure and composition. A large body of literature comparing qMRI markers with histology and advanced physical methods has demonstrated that magnetization transfer saturation (MT_sat_)^42–44^ is sensitive to macromolecular tissue content and, in the brain, reflects tissue myelination. The effective transverse relaxation rate (R2*) is a sensitive marker of brain iron content in addition to myelin^45,46^. A key advantage of qMRI is its quantitative nature and its ability to map the whole brain at high resolution in three dimensions, enabling the characterization of layer-specific myelin profiles. This allows for the generation of non-invasive, whole-brain maps approximating cortical myeloarchitecture across large populations.

In the present study we quantified the microstructure of the hand knob in humans and chimpanzees leveraging the power of ultra-high-resolution qMRI at 7T to compare myelination and iron content^45^ in M1 across species^42,43^. By using qMRI measures of transverse relaxation rate (R2*) and magnetization transfer saturation (MT_sat_)^43^, we characterized tissue iron and myelin content across cortical layers, capitalizing on the link between the qMRI measures and cortical myeloarchitecture (Fig. 1)^42,45,47–50^. We demonstrated that the cortical fields associated with hand control exhibit higher myelination and iron content as compared to those representing other body parts in both humans and chimpanzees. Moreover, lateralization in the microstructure of these fields is stronger in the human motor cortex. These results advance our understanding of the neural architecture underpinning human hand dexterity and handedness, and point to the evolutionary reorganization of motor cortical structures that support the remarkable manual capabilities of our species.

## Results

We used quantitative MRI to map myelination and iron across motor cortical fields in chimpanzees and humans, testing effects of hemisphere, handedness, and age.

### Quantitative comparison of cortical myelin and iron content

We analysed post-mortem, formalin-fixed brains of 16 chimpanzees (8 females, age = 31 ± 19 years [mean ± standard deviation]) with ages covering the chimpanzee lifespan (range = 1.6 to 56 years). The brains were collected using an ethical tissue collection pipeline within the <u>Evolution of Brain Connectomics project (EBC)</u> ^51,52^, see Table 1 for subject details. Microstructural metrics were obtained using quantitative MRI with a multi-parametric mapping protocol providing 300 μm isotropic high-resolution images^47,53,54^. For the between-species comparison, we utilized in vivo multi-parameter maps of 10 human subjects acquired at 500 μm resolution (6 f, [mean age ± SD: 28.0 ± 3.6 y])^55^ on a 7T scanner and 44 human subjects acquired at 800 μm resolution on a 3T scanner (23 f, [mean age ± SD: 28.0 ± 4.1 y]). The analysis of two independent human cohorts acquired at different static magnetic field strengths was used to increase statistical power (using the 3T cohort with larger sample size) while also excluding the potential confounding influence of magnetic field strength when comparing chimpanzees and humans.

**Table 1.** Chimpanzee subject characteristics. Identifier, age (*exact date of birth is known, otherwise: ±1 year for individuals under 10 years old, ±3 years for individuals between 10 and 15 years of age, and ±5 years for individuals > 20 years old ^11^), sex (f = female, m = male), chimpanzee subs^14^cies, acquisition site ^82^d postmortem interval (PMI) are provided.

| Subjects | Age (years) | Sex | Subspecies | Site | PMI (hrs) |
| --- | --- | --- | --- | --- | --- |
| Lukule | 1.6* | m | verus | Sanctuary | <16 |
| Flag | 1.75* | m | verus | Field site | 24 |
| Rasta | 2.75* | m | verus | Field site | 3.5 |
| Emma | 6* | f | verus | Field site | 4 |
| Hoima2 | 13 | f | schweinfurthii | Field site | 15 |
| Hoima1 | 30 | m | schweinfurthii | Field site | 12 |
| Groat | 34* | m | verus | Zoo | 4-6 |
| Rwenzori | 40 | f | verus | Field site | <24 |
| Dorien | 41 | f | verus | Zoo | 4 |
| Tojo | 43 | f | schweinfurthii | Zoo | 1 |
| Rosie | 44* | f | hybrid | Zoo | 1 |
| Fredy | 45 | m | verus | Field site | <18 |
| Jimmy | 45 | m | hybrid | Zoo | <16 |
| Negra | 47* | f | verus | Zoo | 3.5-4 |
| Wilson | 52* | m | verus | Zoo | <24 |
| Sumatra | 56 | f | verus | Field site | 2.5 |

Quantitative MRI parameters for chimpanzees and humans were sampled along the cortical surfaces at ten equidistant cortical depths from the pial surface to the gray/white matter interface yielding laminar markers of cortical myelin (MTsat and R2*), and iron (R2*) content (Figure 1).

We identified the cortical fields of M1 (Brodmann area 4) involved in motor control for foot, hand and face based on anatomical landmarks for both species (Figure 1). Comparison with gold-standard histological delineation of BA4 using Nissl staining in one chimpanzee brain validated our delineation approach in chimpanzee brains (see Supplementary Material-S1).

The average values for both MTsat and R2* in each of the three cortical fields were computed for each individual, each cortical depth providing laminar microstructural metrics for motor cortex subdivisions. For each individual these metrics were compared between cortical fields, pooling across both hemispheres and cortical depth. Additionally, the hand motor fields between the left and right hemispheres were compared to test for lateralization.

### Microstructural characteristics of cortical motor fields

Layer profiles of MTsat and R2* revealed a monotonic increase from the pial surface to the gray–white matter boundary, reflecting known higher myelination and iron content in the deeper cortical layers and an inflection point at medium depths (Figure 2), thereby demonstrating the plausibility of qMRI measures

**Figure 2.**
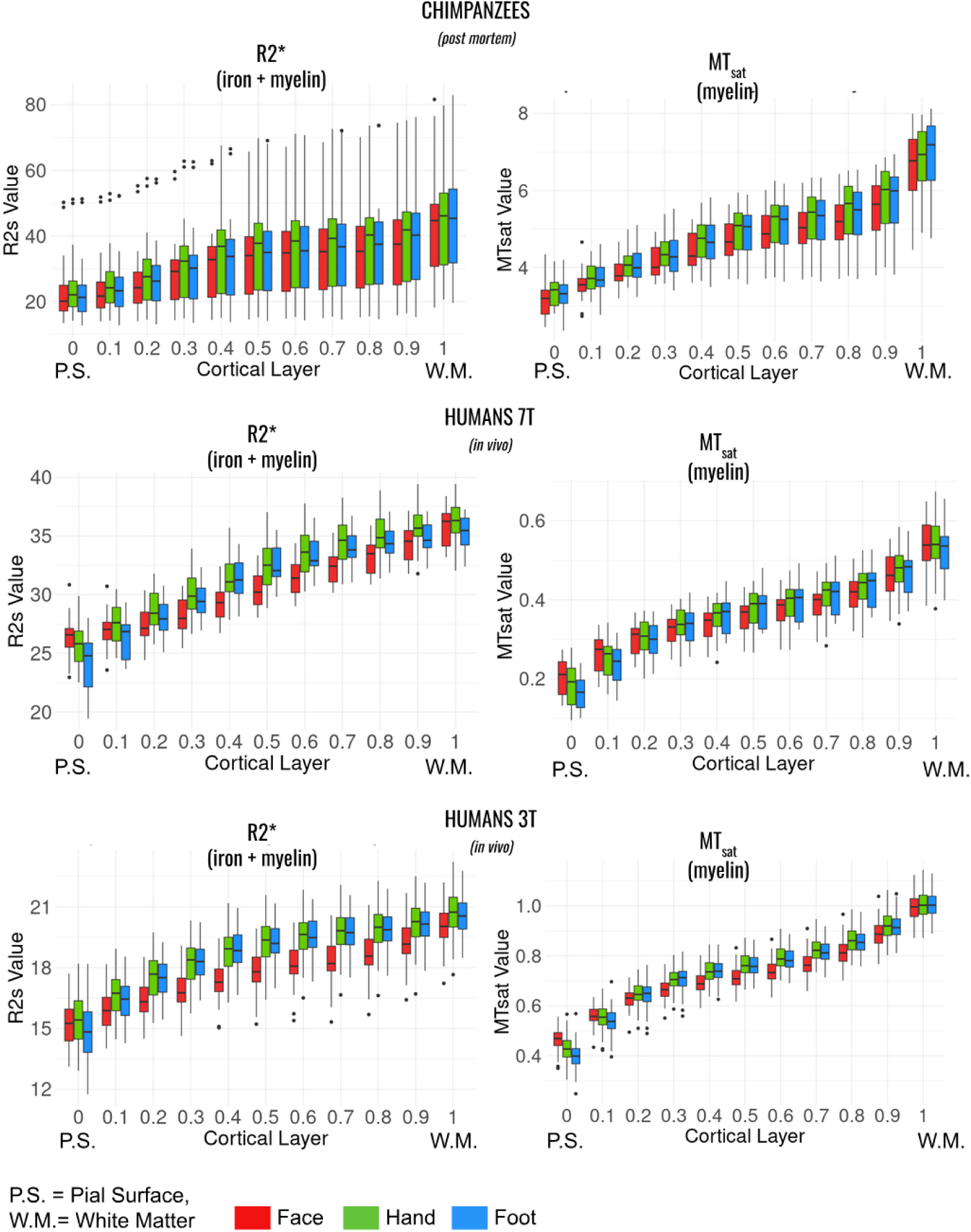
Cortical layer profiles of R2* and MTsat values for the different cortical fields in the human/chimpanzee. Boxplot showing the distribution of R2* (left) and MTsat (right) values across cortical depths for three cortical fields: face (red), hand (green), foot (blue) for 16 chimpanzee brains scanned at 7T (top), and two different cohorts of 10 and 44 human participants scanned at different field strengths (7T and 3T, middle and bottom raws). Boxes = 25/75% percentile; lines = median; whisker = most extreme data value. Cortical depth increases along the x-axis from the pial surface to the white matter boundary. P.M. = Pial Surface, W.M. = White Matter.

We characterized the lifespan trajectories of MTsat and R2* in the chimpanzee dataset spanning the entire chimpanzee lifespan (Supplementary Material-S2). Both MTsat and R2* increased from childhood to adulthood, with characteristic time constants of between 1 and 3 years for MTsat and between 20 and 30 years for R2* years reflecting developmental myelination and age-related iron accumulation respectively. These time constants are in agreement with known literature on humans reporting developmental myelination in first years of life and life-long iron accumulation in the brain^56^.

Differences in the foot, hand and face fields of M1 were observed in both species and across both human cohorts (pooling across hemispheres), with significantly higher R2* in the hand fields than in the foot and the face fields, pointing towards higher levels of cortical iron and myelin in the former (all p<0.01, see Figure 3 and Table 2). Lower R2* values were consistently observed for the face compared to the foot field. Indeed, in chimpanzees, R2* measures were higher by 8.3% compared to face and 3.3% compared to foot. In the human cohort scanned at 7T, R2* measures were higher by 4.8% compared to face and 2.3% compared to foot. In the human cohort scanned at 3T, R2* measures were higher by 6.3% compared to face and 0.9% compared to foot.

**Figure 3.**
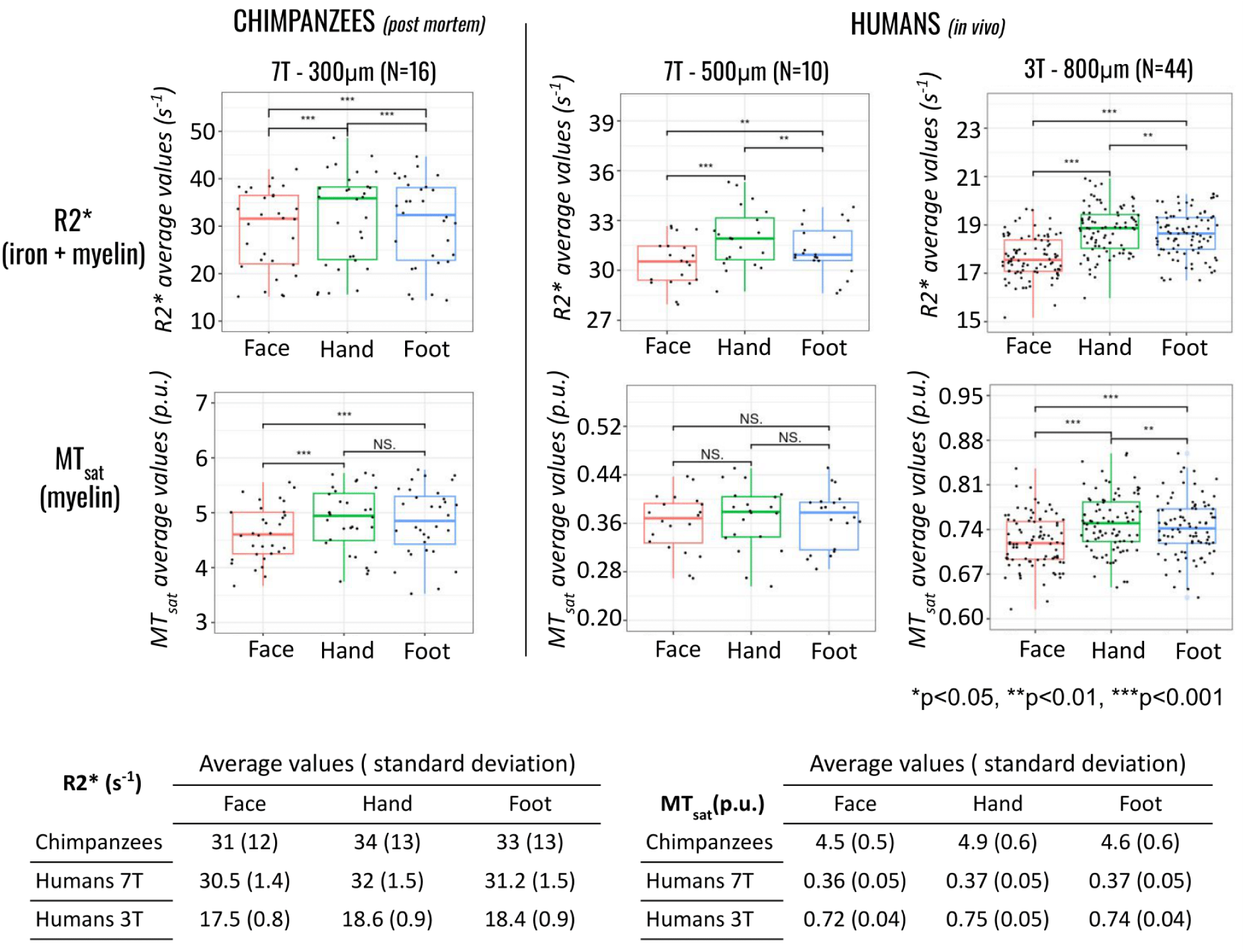
R2* and MTsat values averaged across face, hand or foot motor cortical fields in the human/chimpanzee. Left: Boxplots of the R2* (top row) and MTsat values (middle row) from the cohort of 16 chimpanzee brains scanned at 7T. Right: Boxplots of the R2* (top row) and MTsat (middle row) values from two different cohorts of 10 and 44 human participants scanned at different field strengths (7T and 3T). Bottom row: table of corresponding values for R2* and MTsat maps. Colors: red, green, blue respectively for face, hands and foot fields. Differences in values were examined using paired t-tests. Note the truncated y-axes. Boxes = 25/75% percentile; lines = median; whisker = most extreme data value. *p<0.05, **p<0.01, ***p<0.001 (not corrected).

**Table 2.** Statistical test results comparing qMRI metrics between the different cortical fields in humans and chimpanzees. Corrections for multiple comparisons were made using the Bonferroni correction (p-corr).

|  |  | CHIMPANZEE 7T |  |  |  | HUMAN 7T |  |  |  | HUMAN 3T |  |  |  |
| --- | --- | --- | --- | --- | --- | --- | --- | --- | --- | --- | --- | --- | --- |
|  |  | t-value | p-value | p-corr | df | t-value | p-value | p-corr | df | t-value | p-value | p-corr | df |
| **R2\*** | Face vs Hand | -8.2 | $2.8 \cdot 10^{-09}$ | $8.4 \cdot 10^{-09}$ | 31 | -5.8 | $1.5 \cdot 10^{-05}$ | $4.5 \cdot 10^{-05}$ | 19 | -26.3 | $4.0 \cdot 10^{-43}$ | $1.2 \cdot 10^{-42}$ | 87 |
| | Foot vs Hand | -4.5 | $8.3 \cdot 10^{-05}$ | $2.5 \cdot 10^{-04}$ | 31 | -3.8 | $1.3 \cdot 10^{-03}$ | $4.0 \cdot 10^{-03}$ | 19 | -3.3 | $1.6 \cdot 10^{-03}$ | $4.8 \cdot 10^{-03}$ | 87 |
| | Foot vs Face | 6.3 | $5.4 \cdot 10^{-07}$ | $1.6 \cdot 10^{-06}$ | 31 | 3.3 | $3.4 \cdot 10^{-03}$ | $1.0 \cdot 10^{-02}$ | 19 | 17.1 | $1.6 \cdot 10^{-29}$ | $4.9 \cdot 10^{-29}$ | 87 |
| **MT sat** | Face vs Hand | -6.7 | $1.6 \cdot 10^{-07}$ | $4.9 \cdot 10^{-07}$ | 31 | -1.3 | $2.0 \cdot 10^{-01}$ | $5.9 \cdot 10^{-01}$ | 19 | -11 | $1.6 \cdot 10^{-19}$ | $4.8 \cdot 10^{-19}$ | 87 |
| | Foot vs Hand | -1.1 | $2.6 \cdot 10^{-01}$ | $7.9 \cdot 10^{-01}$ | 31 | -0.4 | $7.3 \cdot 10^{-01}$ | 1.0 | 19 | -2.8 | $5.6 \cdot 10^{-3}$ | $1.7 \cdot 10^{-2}$ | 87 |
| | Foot vs Face | 5.2 | $1.2 \cdot 10^{-05}$ | 3.6 | 31 | 0.9 | $3.8 \cdot 10^{-01}$ | 1.0 | 19 | 6.5 | $4.1 \cdot 10^{-9}$ | $1.2 \cdot 10^{-8}$ | 87 |

In chimpanzees, the MTsat values were significantly lower in the face field than in the hand (by 5.0%) and foot fields (by 4.4%) (p<0.001) indicating higher myelination of the hand motor cortex. We observed 0.6% higher MTsat values for hand than foot fields in chimpanzees, but this difference was not significant. In the large human cohort measured at 3T, MTsat values in the hand field were 4.1% higher than in the face and 0.9 % higher than in the foot fields (p<0.01). A similar trend (non-significant) was observed for the human 7T cohort.

In sum, in both humans and chimpanzees the same pattern of qMRI values emerged – though only partially significant. For both R2* and MTsat the highest values reflecting higher cortical myelin and iron content were observed in the hand field followed by lower values in the foot field, and even lower values in the face field. To further validate the anatomical delineation of motor cortical fields, we repeated the analyses using independent functional motor activation maps derived from the Human Connectome Project and restricted to primary motor cortex (BA4). These analyses yielded highly consistent results across both human cohorts, with the hand representation exhibiting significantly higher R2* and MTsat values compared to face/tongue and foot representations (Supplementary Material - S3 and figure S3). This confirms that the observed microstructural differences across motor fields are robust to the method used for region definition.

### Lateralization of the hand field

In chimpanzees, no significant differences between the left and the right hemispheres in the hand field were observed for R2* and MTsat values (Figure 4).

**Figure 4.**
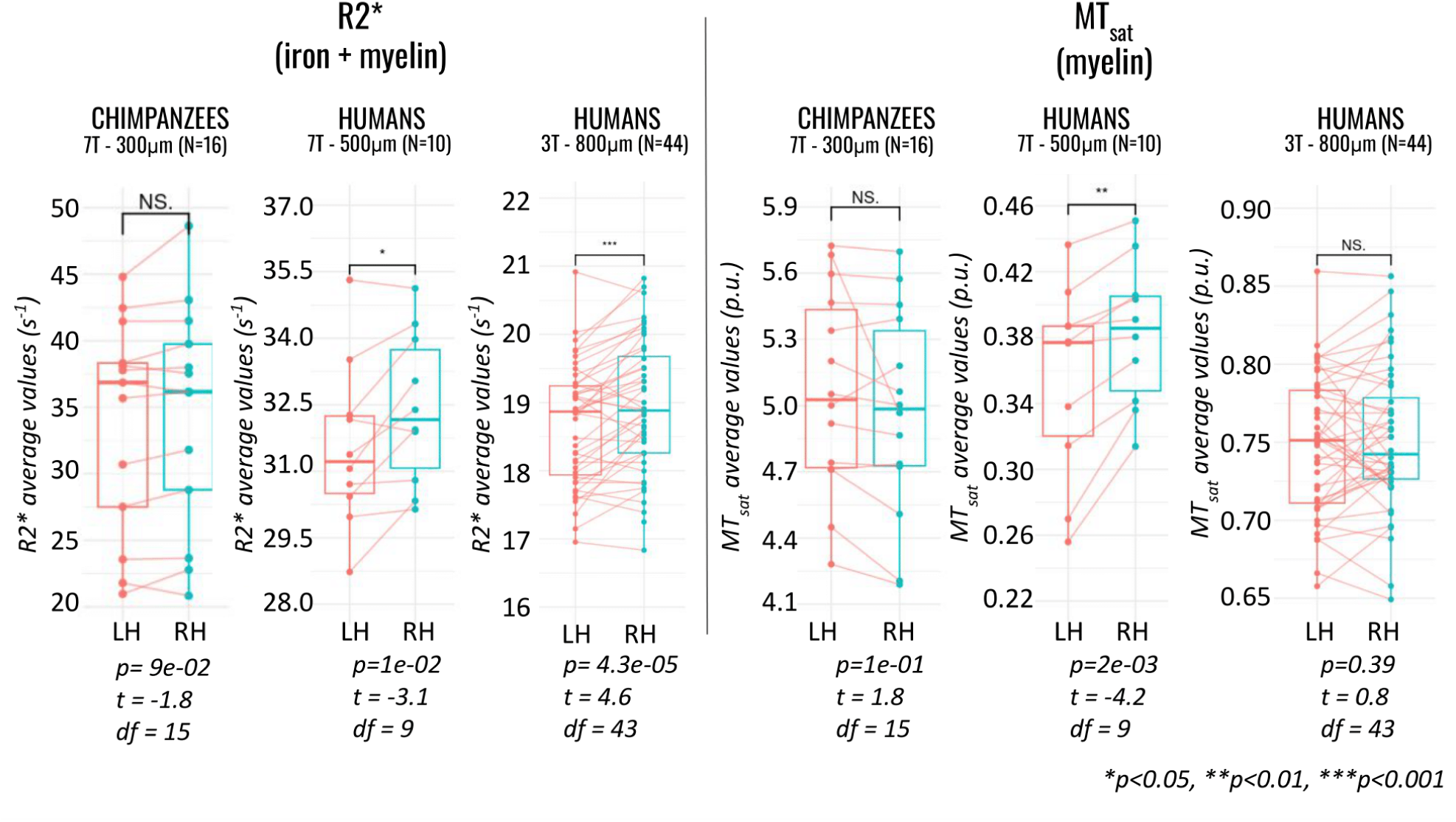
R2* and MTsat averaged across the hand motor cortical field for left/right hemispheres and human/chimpanzee. Boxes = 25/75% percentile; lines = median; whisker = most extreme data value excluding outliers. Left: Boxplots of R2* values in cohorts of chimpanzee and humans (7T and 3T); Right: Boxplots of MTsat values in cohorts of chimpanzees and humans (7T and 3T). Colors: red for the left hemisphere and blue for the right hemisphere hand cortical fields. Datapoints of the same individuals are linked by a line. Note the truncated y-axes. Differences in values were examined using paired t-tests, *p<0.05, **p<0.01, ***p<0.001.

In humans, R2* values were higher in the right hemispheric hand field than in the left for both the 3T (by 1.8%, p<0.001, t=4.6, df=43) and 7T (by 2.7%, p<0.05, t=3.1, df=9) cohorts. MTsat values were significantly higher (by 7.7%) in the right than in the left hemisphere hand field in the 7T cohort (p<0.01, t=4.2, df=9) but not in the 3T cohort.

In comparing asymmetry quotients (AQ=2x[values_left_-values_right_]/[values_left_+values_right_]) in R2* between the different species, we observed significant differences between chimpanzees and the two human cohorts (Fig. 5, all p<0.05) with humans showing higher lateralization (t=2.1, p=0.02 testing for chimpanzees and the human cohort scanned at 7T and t=1.8, p=0.04 testing for chimpanzees and the human cohort scanned at 3T), but no significant differences between the two human cohorts (t=-0.9, p=0.82), see Figure 5, top row.

**Figure 5.**
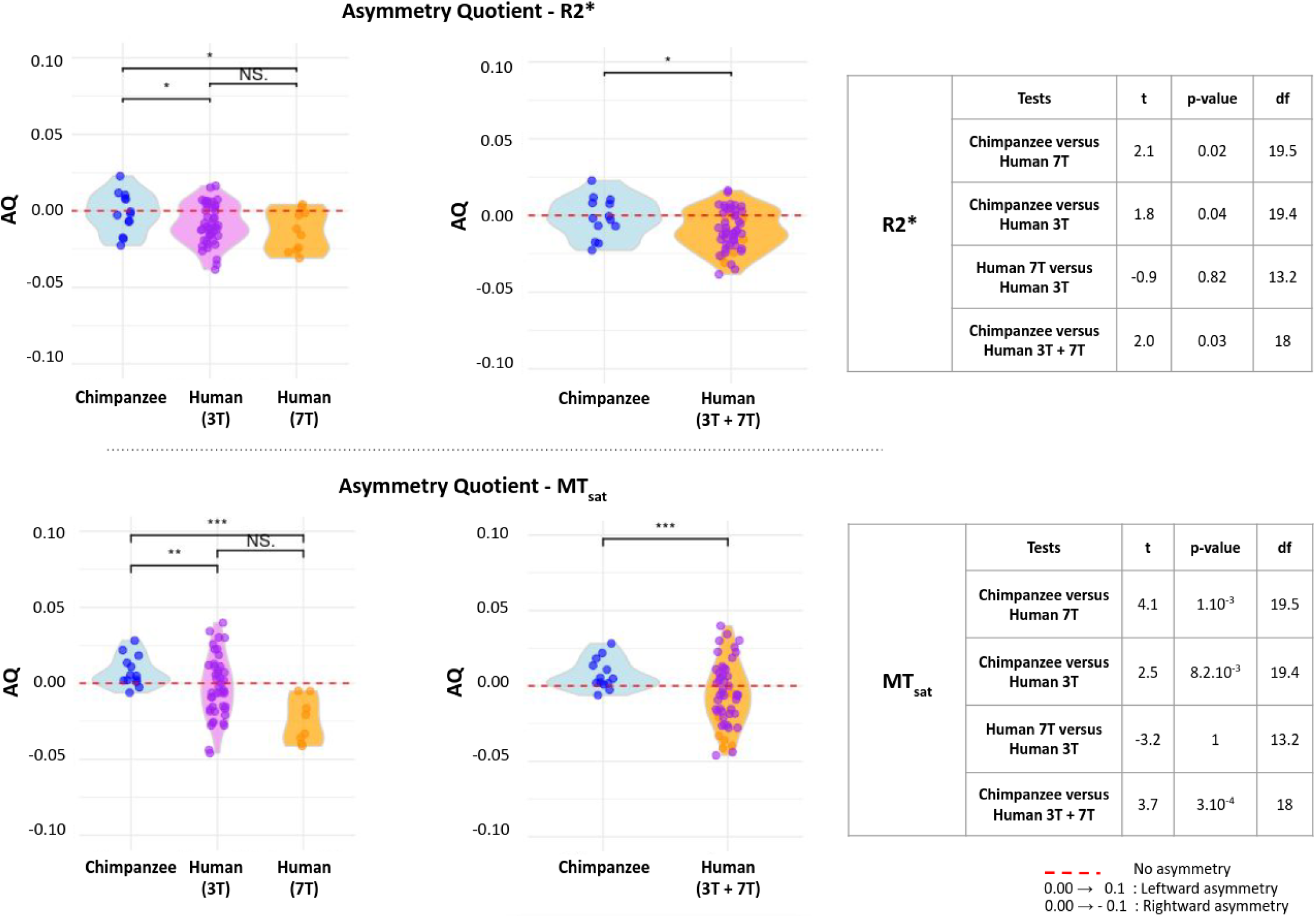
Distribution of R2* and MTsat asymmetry quotients for chimpanzees and humans. The violin plots display the distributions of asymmetry quotients of R2* and MTsat values for chimpanzees and humans (7T and 3T cohorts separately [left panels] and pooled [right panels]). Individual data points are shown as jittered dots, with colors indicating the specific groups: blue for chimpanzees, dark orange for humans scanned at 7T, and purple for humans scanned at 3T. The dashed red line represents an asymmetry quotient of zero. T-tests were performed to compare the mean MTsat and R2* asymmetry quotients between chimpanzees and the combined human group. The results of the t-tests (two-sample, unpaired t-test with unequal variances - Welch’s t-test) are listed next to each plot.

Similar results were obtained for asymmetry quotients using MTsat with significant differences between chimpanzees and both human cohorts (both p<0.01) with higher lateralization in humans (t=4.1, p=0.001 testing for differences of chimpanzees and the human cohort scanned at 7T and t=2.5, p=0.008 testing for differences of chimpanzees and the human cohort scanned at 3T), but no difference between the two cohorts of human participants (t=-3.2, p=1), see Figure 5, bottom row. Using the functionally defined hand motor regions restricted to BA4, hemispheric asymmetries in humans were similarly observed and remained consistent with the main findings (Supplementary Figure S4), indicating that lateralization effects are preserved when independent functional definitions of the hand motor field are applied. To assess the potential influence of age distribution in the chimpanzee cohort, we repeated the analyses after excluding individuals younger than 6 years. The pattern of microstructural differences across motor cortical fields remained unchanged, with higher R2* and MTsat values in the hand representation (Supplementary Figure S5). However, hemispheric asymmetry in the hand field remained non-significant in this restricted sample (Supplementary Figure S6), consistent with the main analysis.

## Discussion

This study examined interspecies differences and similarities in microstructural characteristics of M1, specifically in regions controlling hand movements in humans and chimpanzees using high-resolution quantitative MRI. Surprisingly, the microstructure of the motor cortex related to its somatotopic organization is relatively unexplored and controversial in human and non-human primates. Unique access to chimpanzee brains, ethically sourced through the international EBC project, allowed us to explore this question in this scarce and endangered species.

Previous studies in humans examining the structure of the sensorimotor system gave insights into possible brain asymmetry and handedness and sparked debate in the field. Using quantitative histological and imaging techniques, White et al.^57^ found no asymmetry of S1, while earlier work by White et al.^58^ identified cerebral asymmetry correlating with handedness. In particular, their results suggested that humans have more cortical circuitry devoted to the representation of the right upper extremity than to the left. Consistent left hemisphere dominance in neuropil volume in various cortical areas in humans was observed by Amunts et al.^37^. Similarly, Foundas et al.^59^ observed asymmetries in the central sulcus linked to hand preference, revealing a significant leftward asymmetry in the motor hand area, larger in the precentral gyrus in right-handers, but no consistent directional asymmetry in left-handers.

Concerning the chimpanzee motor cortex microstructure, histological studies from Sherwood et al.^11^ showed that neuron density in layer II/III of M1 was significantly higher in the left handknob area, but no population-level asymmetry was observed in other histological features, contrasting with human studies that have shown a consistent left hemisphere dominance in neuropil volume^37^. This suggests that the relationship between asymmetry measures of cellular volumes of M1 and handedness in humans, at the population level, may be a unique evolutionary trait. Although studies on macroscopic anatomy and some histological studies of the chimpanzee’s motor cortex are available, to our knowledge no histological studies on the myelin content in the different cortical fields or lateralization of myelination in the motor cortex were published.

We leveraged the quantitative capabilities of qMRI, which provides indirect estimates of myelin and iron content that are comparable across brain regions, between hemispheres, and even across species, all within large sample sizes. Such comprehensive comparisons are difficult to achieve with histological studies, which typically offer only qualitative measures of myelin and iron in limited tissue sections and are usually based on small numbers of individuals.

By combining the unique brain resource with advanced neuroimaging methods, this study adds to the literature by showing that the different motor fields exhibit different tissue properties in both species, and that only humans show population-level hemispheric asymmetry in the hand-related motor cortex.

In both humans and chimpanzees, the hand field of M1 exhibited higher MTsat and R2* values than the face and foot fields, consistent with prior findings in humans^31^, which demonstrated higher myelination in the hand compared to the face and highlighted a similarity shared with our closest living primate relatives. Since myelin supports fast and efficient conduction but may also reduce neural plasticity, these results allow us to speculate about a strong predisposition towards hand-related behaviors, flexibility, and strength in our common ancestor from whom we diverged between 7 and 8 million years ago^60^.

Both human and chimpanzee M1 exhibited a similar pattern of iron and myelin content, with highest values in the hand field, followed by the foot, and the face cortical fields. Importantly, this pattern was replicated when motor cortical regions were defined using independent functional activation maps restricted to primary motor cortex, supporting the robustness of the observed microstructural organization across methods of region definition. This shared pattern may reflect ancestral adaptation linked to object manipulation and locomotion. However, species differences in usage and motor repertoire likely play a role. While humans perform a broad range of skilled, bimanual tasks, chimpanzees also engage in complex manipulations (e.g., tool-use, social gestures) but also quadrupedal locomotion^2^.

When interpreting qMRI results, it is important to recognize that cortical myelin and iron content detected by qMRI is an integral measure of both radial fibers that connect the cortex to distant areas and tangential intracortical short-range fibers that support local computation. Therefore, different levels of iron and myelin in motor cortex subdivisions and their lateralisation may be driven by differences in the density, myelination rate and iron content of one or both fibre populations.

This study determined for the first time potential microstructural correlates of handedness in the human motor cortex using non-invasive high-resolution qMRI. Typically, behavioral asymmetries are believed to be reflected in the neural specialization in the contralateral brain hemisphere. This is largely due to the partial or complete crossing of the majority of body-brain connections, in case of the motor system crossing over primarily at the pyramidal decussation in the medulla oblongata. Thus, motor control is predominantly exerted by the opposite hemisphere of the brain. In other words, the left hemisphere controls the right hand and vice versa. Surprisingly, the human cohorts studied, composed of right-handed volunteers, showed higher myelin- and iron-sensitive qMRI parameters in the right hemisphere hand field. This challenges a common conjecture that the left motor cortex in right-handed individuals should be more highly myelinated, which has not been histologically demonstrated. One interpretation is that dominant-hemisphere motor control may require more plasticity and therefore retain lower myelination levels of intracortical fibers^61^. Since it has been reported that motor skill learning is faster for the preferred hand^62^, and that this preference develops very early in infancy^63^, we could reasonably speculate that lower myelination of the left hemisphere cortical field may be kept to facilitate plasticity in learning new motor sequences for the right hand.

At the population level, the lateralization effect did not reach significance in chimpanzees. Additional analyses accounting for age distribution by excluding younger individuals did not alter this conclusion, as hemispheric asymmetry remained non-significant in the chimpanzee cohort. This suggests that the absence of population-level lateralization in chimpanzees is unlikely to be solely driven by developmental variability within the sample. However, notable hemispheric differences were detected at the individual level, with some chimpanzee subjects exhibiting higher values in either the left or right hemisphere. This is consistent with previous studies on laterality in chimpanzees, indicating that heterogeneity in lateralization may be higher in chimpanzees than in humans^8^, masking lateralization effects at the population level. Nevertheless, individual hand preferences appear to exist based on field observations^4^ as well as in our measurements. However, we could not investigate the relation between motor behavior and brain structure at the level of the individual chimpanzee due to the lack of individual motor behavior assessments.

The stronger lateralization in humans compared to chimpanzees could stem from the unique fine movement capabilities and dexterity that humans developed during evolution and in infancy^64^. Additionally, humans are strongly encouraged to use their right hand from an early age. This early training coincides with phases of substantial brain development and may coincide with enhanced plasticity in the motor cortex^65^. It is reasonable to acknowledge that we cannot expect the same level of dexterity from chimpanzees due to differences in muscle anatomy. Humans possess a robustly muscular and fully opposable thumb. Conversely, chimpanzees possess a smaller opposable thumb compared to humans, with a muscle arrangement that allows for less strength and mobility, consequently offering limited precision and flexibility in finger movements^66^.

### Considerations and limitations

Due to the scarcity of functional information for the studied chimpanzee brains and the lack of a species-specific cortical atlas, our somatotopic labeling relied on anatomical landmarks. We utilized the hand-knob as a reliable sulcal marker for dorso-ventral boundary, but delineation of the hand, face, and foot regions may have included adjacent motor areas. While we carefully referenced human functional and histological boundaries (BA4–6, BA3–4 ^67^), and confirmed the location of BA4 using Nissl-stained histological sections in one individual^68^ (see Supplementary Material-1), finer mapping, especially for the subdivision of the face field containing the mouth and tongue areas, remains challenging in the absence of recent fMRI data in chimpanzees.

The utilised qMRI parameters R2* and MTsat have previously been shown to relate to iron content and myelination (for an overview, see (Weiskopf et al., 2021^43^)). The use of qMRI parameters is further supported by the similarity of whole brain qMRI maps and known regional variations in myeloarchitecture (Fig. 1), the comparable motor cortex delineation using the qMRI maps and gold-standard histology and the similarity of qMRI cortical depth profiles and myeloarchitectural profiles (Fig. 2). However, these qMRI markers only indirectly reflect these microstructural features and are not exclusively influenced by them. Thus, we cannot exclude the possibility that results were affected by other microstructural features and findings should be interpreted with caution and not equated with direct measurements of cyto- or myeloarchitecture.

Although hand use was measured across several of the chimpanzee populations studied, there were too few individuals in the sample for which there are both hand preference and post-mortem brain measures. Studies suggest that rather than a species-level hand preference, individuals may either demonstrate left, right or no hand preference, hence assessment at the individual level is needed to assess an relevant brain structural asymmetry. Consequently, we were not able to directly relate hemispheric microstructural asymmetries to individual hand preference. This limitation should be considered when interpreting inter-individual variability in lateralization.

## Conclusion

This study showed differences and similarities in the microstructural features across the motor cortex of chimpanzees and humans.

Levels of qMRI-estimated myelin and iron content were higher in the hand-knob area than in areas representing other body areas in both human and chimpanzee species, pointing to an evolutionary conserved trait. An important species difference was the stronger lateralization of myelination and iron content in the hand-knob of humans than in chimpanzees, in line with more pronounced handedness of humans when performing fine hand and finger movements. This implies that the evolution of hand motor skills involves not only alterations to sulcal anatomy but also significant changes in cortical microstructure. These findings provide valuable insights into the anatomical underpinnings of distinct motor behaviors, such as extensive tool-use observed in hominins.

## Materials and Methods

### Chimpanzee and human cohorts

#### Chimpanzee subjects

The x were obtained following the standard operating procedures of the EBC project^51,52^. The strengths of this project lie in its ethical methodology and multi-disciplinary methods. When chimpanzee individuals naturally die either in the wild or within sanctuaries/zoo environments, the researchers are afforded the opportunity to gain access to their brains for investigative purposes.

Brains from wild and captive chimpanzees (Pan troglodytes) were sourced from African field sites, zoos and sanctuaries. The causes of death for chimpanzees from field sites included bacterial infections (N = 1), human-animal conflict (N = 2), leopard attack (N = 2), starvation (N=1), chronic kidney disease (N=1), intergroup attack (N=1). The cause of death for chimpanzees that died at sanctuaries and zoos were conspecific aggression (N = 1), renal disease (N = 1), pneumonia (N=1), nephritis (N=1), stroke (N=1), cardio-vascular diseases (N = 2), epileptic episode (N = 1).

The brains were extracted within 4-24 hours after death and fixed for 8-12 weeks in phosphate-buffered saline (PBS) with 4 % of paraformaldehyde. The samples were then shipped to Leipzig where superficial blood vessels were removed and the formalin was washed out in PBS with 0.1 % sodium azide at pH 7.4. The procedures were in line with the ethical guidelines of primatological research at the Max Planck Institute for Evolutionary Anthropology, Leipzig, which were approved by the ethics committee of the Max Planck Society. Sixteen chimpanzee brains (8 f, age 1.1-52.0y) were included in this study, further details can be found in^42^.

#### Chimpanzee brain MRI scanning

The methodological details of the scanning are described in detail in^47^. We provide here only a brief overview. Chimpanzee brains were placed in acrylic containers filled with proton-free fluid Fomblin and scanned on a human whole-body 7T Terra MRI scanner (Siemens Healthineers, Erlangen, Germany), using a 32-channel receive, single channel transmit radio-frequency (RF) human head coil (Nova Medical, Wilmington, MA). Multi-parametric mapping^53,54^ was implemented using three acquisitions of a multi-echo 3D fast low angle shot (FLASH) sequence at 300 μm isotropic resolution (matrix: 432 x 378 x 288; readout bandwidth of 331 Hz/pixel; repetition time (TR) = 70 ms; 12 equidistant echoes with echo times (TE) between 3.63 ms and 41.7 ms with a bipolar readout, ΔTE = 3.56 ms; excitation flip angles: 18° (for PD-weighted and MT-weighted acquisition), 84° (for T1-weighted acquisition). For magnetization transfer weighting a Gaussian pulse at 3 kHz offset with nominal flip angle of 700° was used.

Maps of the RF receive bias field (B1-) were estimated using the ratio between a low resolution (2.1 mm) T1-weighted image acquired with the 32-channel receive coil and a second low resolution T1-weighted image acquired with the bird-cage transmit element of the head coil in receive mode and used to correct the quantitative maps for receive bias^69^.

Maps of the RF transmit field B1+ were measured using a spin and stimulated echo measurement sequence in combination with a dual gradient echo sequence that mapped the static magnetic field B0^70^. B1+ mapping was done at an isotropic resolution of 4 mm, using a 3D echo-planar imaging (EPI) readout the following parameters: TR = 500 ms; TE = 40.54 ms; mixing time TM = 34.91 ms; flip angles 330° to 120° in steps of 15°; RF duration 24 μs per °; generalized autocalibrating partial parallel acquisition (GRAPPA) acceleration factor = 2 x 2. B0 mapping was done using a dual-echo FLASH acquisition with an isotropic resolution of 2 mm, using TR = 1020 ms, TE = 10 and 11.02 ms, flip angle = 20°. The B0 map was used to correct the EPI distortions in the B1+ map.

#### Human brain MRI scanning

Quantitative multi-parameter mapping in vivo datasets of different published human cohort studies were used for comparison with chimpanzee data. All volunteers were right-handed.

The first dataset consisted of 7T high resolution qMRI data acquired at 500 μm of ten healthy adult human right-handed participants (6 f, [mean age ± SD: 28.0 ± 3.6])^55^. All subjects gave written informed consent and the study was approved by the ethics committee of the medical faculty of the University of Leipzig (Reg.-No. 273-14-25082014). As presented in^55^, the MPM protocol involved three multi-echo FLASH scans with T1- and PD-weighting (T1w, PDw, and MTw), along with maps of the RF transmit field B1+ and the static magnetic field B0. The MPM acquisition had whole-brain coverage at an isotropic resolution of 500 μm. The PD-weighted and T1-weighted multi-echo FLASH scans were acquired with flip angles of 5° and 24°, respectively. Readouts with alternating polarity were used to produce six evenly spaced echoes between 2.8 and 16 ms, with a TR of 25 ms, resulting in a total imaging time of 18 minutes per volume. Additional parameters included a matrix size of 496 × 434 × 352 (read × phase × partition), sagittal orientation, GRAPPA with an acceleration factor of 2 in both phase and partition directions (inner phase encoding loop), non-selective excitation with a sinc-shaped RF pulse, and a readout bandwidth of 420 Hz/pixel. The transmit voltage was optimized for the occipital lobe using an initial low-resolution transmit field map. Motion was monitored and corrected prospectively by an optical tracking system (Kineticor, Honolulu, HI). For the prospective motion correction of the high-resolution MPM acquisitions, each participant was scanned while wearing a mouthguard assembly with attached passive Moiré pattern markers, molded to their front teeth (manufactured by the Department of Cardiology, Endodontology, and Periodontology, University Medical Center Leipzig).

The second dataset consisted of 44 adult right-handed participants (23 f, [mean age ± SD: 28.0 ± 4.1]); the study was presented to and approved by the ethics committee of the medical faculty of the University of Leipzig (ID: 293/18-ek). The subjects were scanned using a 3T Connectom scanner equipped with a 32-channel receive RF head coil (Siemens Healthineers, Erlangen, Germany) and a body transmit coil. Multi-parametric mapping was implemented using a multi-echo 3D FLASH sequence at 800 μm isotropic resolution (matrix: 320 x 280 x 224; readout bandwidth of 488 Hz/pixel; TR = 25 ms; 8 equidistant echoes for PD-weighted and T1-weighted images and 6 equidistant echoes for MT-weighted images, with TEs between 2.4ms and 18.64 ms (14 ms for MT-weighted), △TE = 2.32 ms; excitation flip angles: 6° (PD-weighted and MT-weighted), 21° (T1-weighted); MT pulse characteristics: Gaussian at 2 kHz offset, nominal flip angle: 180°, 4 ms. Additional parameters: GRAPPA acceleration factor = 2 × 2. The acquisition time for each of the weighted images was about 7 min. B1+ mapping was done at an isotropic resolution of 4 mm, with a matrix size of 64 x 64 x 48 and the following acquisition parameters: TR = 500 ms; TE = 43.5 ms; mixing time TM = 38.2 ms; flip angles 230° to 130° in steps of 10°; RF duration 28 μs per °; GRAPPA acceleration factor = 2 x 2 ^71^. B0 mapping to correct the EPI distortions of the B1+ maps was done at a resolution of 3 x 3 x 2 mm and a matrix size of 64 x 64 x 64, with the following acquisition parameters: TR = 1020 ms, TE = 10 and 12.46 ms, flip angle = 90°. The RF receive bias (B1-) field was partly corrected using the ratio between the low resolution (4 mm) PD-weighted image (excitation flip angle 6°, TR 6 ms, TE 2.4 ms) acquired with the 32-channel RF receive coil and an another similar image acquired with the body RF coil in receive mode. Separate receive bias field maps were acquired for the PD-, T1- and MT-weighted acquisitions to account for the inter-scan motion effects on the bias field^69^.

#### Creation of qMRI maps for postmortem chimpanzee brains

The maps of transmit RF field B1+ were generated using the open-source hMRI toolbox (https://hmri.info) assuming a global reference T1 of 500 ms, accounting for the shortening of T1 in formalin-fixed tissue. A boundary-preserving smoothing procedure ensured sharp edges between the brain and the no-signal background. A similar processing strategy was employed for the receive B1- maps, calculated by dividing the image intensity from the 32-channel receive coil by the acquisition with the transmit coil in receive mode (averaged over the first five echoes).

All weighted 3D GRE FLASH images were corrected for susceptibility-induced distortions, using odd and even echoes recorded with the opposite readout polarity and the HySCO algorithm from the ACID toolbox (http://diffusiontools.com/). The effective transverse relaxation rate (R2*) was fitted using weighted least squares exponential fit as implemented in the hMRI toolbox. From this fit, the weighted images were extrapolated to TE = 0, and these images were used to calculate R1, PD, and MTsat as described previously. The MTsat images were further corrected for residual B1+ bias using the algorithm described in^47^.

#### Creation of qMRI maps for human brains

B1⁺ maps were estimated using a global reference T₁ value of 1192 ms, the default in vivo 3T setting in the hMRI toolbox. For the post-mortem chimpanzee MRI data, the effective transverse relaxation rate (R2) was computed using weighted least squares fitting, and the weighted images were extrapolated to TE = 0. These extrapolated images were then used to generate MTsat, R1, and PD maps, following the hMRI toolbox pipeline^72^. The processing also included per-contrast RF sensitivity bias correction and a correction for imperfect spoiling.

In both humans and chimpanzee brains, the magnetization transfer saturation (MTsat) map served as a myelin marker^49^ and the effective transverse relaxation rate (R2*) map as a marker for myelin and iron content^45^; see Figure 1, B. We note that they are only indirect measures of myelination and iron content that are also influenced by other tissue components (e.g. hydration levels) but were widely used as such markers^43^.

### Post-processing and segmentation

To segment the chimpanzee brain data, we first created brain masks by thresholding the T1-weighted images acquired at the shortest echo time, since the scans were performed in a proton-free (MR-invisible) solution. For optimal gray-white matter contrast, MTsat images were downsampled to 0.7 mm isotropic resolution. We then used the Freesurfer pipeline (https://surfer.nmr.mgh.harvard.edu) for cortical segmentation and surface reconstruction. The process included 1) running *autorecon1* (with *-noskullstrip -notal-check -hires*), 2) converting our custom brain mask to Freesurfer format using *mri_convert --conform_min,* 3) running autorecon2 (*-notal-check -hires*), 4) Manually editing white matter masks and contrast in *brain.mgz* when needed using Freeview, 5) re-running white matter segmentation with *autorecon2-wm*, 6) refining pial surfaces using *autorecon-pial*. Further details about the data processing pipeline can be found in ^42,47^.

Cortical surfaces for the in vivo human data (3T) were reconstructed using FreeSurfer’s *recon-all* pipeline. Because the contrast in qMRI parameter maps differs substantially from the T1-weighted MPRAGE contrast expected by recon-all, additional preprocessing steps were performed to generate a synthetic image with MPRAGE-like contrast from the 3T qMRI maps. First, estimation errors were corrected by removing a small number of negative or abnormally high values from the R1 and PD maps. These cleaned R1 and PD maps (with T1 = 1/R1) were then used as input to FreeSurfer’s mri_synthesize routine to create a synthetic FLASH image optimized for white/gray matter contrast (TR = 20 ms, FA = 30°, TE = 2.5 ms).

This synthetic image was used as input to *autorecon1* with the *-noskullstrip* flag. Skull stripping was then performed using a brain mask derived from tissue probability maps (GM/WM/CSF, threshold > 0) generated by the SPM Segment tool (https://www.fil.ion.ucl.ac.uk/spm). The resulting skull-stripped image was passed into the remaining stages of the recon-all pipeline.

Cortical surfaces for the in vivo human data (7T) were reconstructed using FreeSurfer’s *recon-all* pipeline (v6.0;^73^). As this pipeline is optimized for standard T1-weighted MPRAGE images, several adaptations were necessary for the 7T quantitative MRI (qMRI) data. A synthetic FLASH image with MPRAGE-like contrast (TR = 20 ms, FA = 30°, TE = 2.5 ms) was generated using FreeSurfer’s *mri_synthesize* tool, with scaled PD and T1 maps (T1 = 1/R1) as inputs. Prior to synthesis, erroneous negative and extremely high values were removed. To improve skull stripping, SPM’s Segment tool was used to create a combined GM/WM/CSF tissue probability mask (threshold > 0), which was applied to the PD map. This masked PD image was normalized so that white matter intensity averaged 69%^72^, then inverted (100% – PD) to resemble MPRAGE contrast. Rician denoising^74^ was applied, and the resulting image was input into recon-all for cortical surface reconstruction.

Further details can be found in McColgan et al. 2021^55^.

#### Cortical labeling

The representations of the different body areas in the motor cortex were approximated based on the anatomical definition of the hand-knob via sulcal structures. The hand representation in M1 (BA4) was manually defined on chimpanzee and human brains’ pial mesh surfaces using the sulco-gyral motif, using the FreeSurfer labeling tool available in Freeview (https://surfer.nmr.mgh.harvard.edu, see Figure 1, A.).

To determine the boundaries between BA4 (M1) and the adjacent areas BA3 (S1) and BA6 (premotor cortex), we used established anatomical landmarks from humans and non-human primates. The posterior boundary of BA4, marking its transition to BA3, was defined by the base of the central sulcus, where the sulcus shifts from the precentral gyrus (motor cortex) to the postcentral gyrus (somatosensory cortex). This transition is characterized by a change in the gyral pattern, visible as a flattening at the floor of the central sulcus.

The anterior boundary of BA4, marking its transition to BA6, was defined by the crest of the precentral sulcus, which separates M1 from the premotor cortex. This boundary placement is supported by converging evidence from previous human^75,76^, non-human primate^77^ and monkey^78^ myeloarchitectonic and cytoarchitectonic studies. These findings guided our delineation of the BA4 and BA6 boundary in both species. We further checked this definition by comparison to the cytoarchitectonic criteria in one post-mortem brain subject, identifying these borders by differences in cortical thickness and cell types (e.g., the granular layer in BA3 and the agranular cortex in BA4) and the presence of Betz cells, essential for motor control^79^ in BA4 (see also Suppl. Material 1).

The dorsal and ventral boundaries of the hand-knob within the central sulcus were identified within the precentral gyrus to demarcate the hand area. Adjacent to the hand cortical region along the precentral gyrus, the dorsal region was designated as the foot/leg area, extending dorsally to the inter-hemispheric fissure and further to the depth of the para-lingual gyrus. Ventral to the hand area along the precentral gyrus, we labeled the oro-facial region (hereafter referred to as the face area for simplicity).

To further validate the anatomical delineation of motor cortical fields, we performed an additional analysis using independent functional motor activation maps from the Human Connectome Project (HCP). Group-level activation maps corresponding to hand, foot, and tongue movements were projected onto the fsaverage surface and converted into cortical labels. Because these functional activations include both motor and somatosensory components, the resulting regions were restricted to the precentral gyrus (BA4) using the same anatomical criteria described above to ensure consistency with the primary motor cortex definition used in the main analyses. These functionally derived ROIs were then applied to the human datasets, and the same microstructural sampling and statistical analyses were performed (see Supplementary Material S3).

### Statistical analyses

#### Comparing R2* and MTsat values averaged across face, hand and foot brain motor cortical fields in the human and chimpanzee species

R2* and MTsat values for each of the cortical fields representing foot, hand and face, labeled for both species, were sampled across the different brain cortical laminae, ranging from the most external pial surface to the gray/white matter interface, encompassing ten computational laminae in total using Matlab (R2021a; The Mathworks, Natick MA). Values were first averaged across all ten cortical depths/laminae and subsequently averaged across all voxels belonging to the same label.

To evaluate possible differences between the different cortical fields within the chimpanzee and human cohorts, paired t-tests were performed between the different fields, pooling the left and the right hemisphere values for each field. Significance values were corrected for multiple comparisons using the Bonferroni correction. The significance threshold was p = 0.05.

#### Hemispheric comparison of R2* and MTsat values within each species

To examine differences between the left and right hemisphere hand cortical fields chimpanzee and human cohorts (7T and 3T), paired t-tests were conducted on R2* and MTsat within each species.

The paired t-test was chosen since data points for the left and right hemispheres were present for each subject, allowing for a direct comparison. The test was two-sided, testing for differences in either direction, and assumed equal variances between the groups. The significance threshold was p = 0.05.

#### Hemispheric comparison of R2* and MTsat values within each species

To quantify the laterality of the hand cortical field, a laterality index was computed for each brain region, for both chimpanzees and humans. The asymmetry quotient (AQ) was defined as the normalized difference between the mean values of the left hemisphere (LH) and the right hemisphere (RH) of the brain. Specifically, for each species, the laterality index was calculated using the formula:

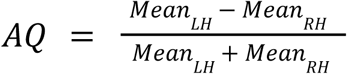

Two sets of comparisons were performed to evaluate differences in laterality between chimpanzees and humans, as well as between the two human cohorts:

– Chimpanzees vs. Humans (7T and 3T separate): A violin plot was created for each species, showing the distribution of AQ for R2* and MTsat. For humans, the data were plotted separately for the 7T and 3T scanner cohorts. Comparisons between chimpanzees, humans scanned at 7T, and humans scanned at 3T were performed using Welch’s t-test, a two-sample unpaired t-test that does not assume equal variances between groups.
– Chimpanzees vs. Humans (7T and 3T combined): the human data scanned at 7T and 3T scanners were pooled into a single group. This was done to assess overall differences between chimpanzees and humans, regardless of the specific scanner used. A Welch’s t-test was performed to compare the pooled human group with the chimpanzees.

The significance threshold was p = 0.05.

#### Exploring the lifespan trajectory of R2* and MTsat values for each cortical field within the chimpanzee species

We implemented a Bayesian model using PyMC (v.5.16.1) in Python to estimate the time constant for two brain imaging measures, R2* and MTsat, across the different cortical fields corresponding to foot, hand and face:

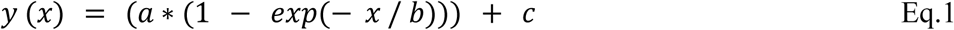

where *y* represents the level of iron or myelin estimated from qMRI parameters at a certain age; *x* represents the age; *a* represents the total increase across the entire lifespan, giving the maximal value reached at old age; *b* represents the characteristic time constant, such that the saturation rate of lifespan changes is 1/b; and *c* represents the constant term, which describes the value at birth. This model was previously employed by^56^ to explore the lifespan trajectory of iron accumulation in the human brain. The time constant here describes how quickly the R2* and MTsat values change over the lifespan in the different cortical fields.

Visualization and diagnostics were conducted using Arviz (v.0.18.0). To ensure reliable estimates, we discarded the first 2000 samples drawn from the model for model tuning and then collected an additional 2000 samples for analysis, across 4 independent chains. We assessed the quality of the model fit by checking for chain convergence using the Ȓ statistic^80^, with detailed results presented in the Supplementary Materials.

Priors and Model Assumptions - We assumed weakly informative prior distributions for the model parameters based on existing knowledge. For each brain region, we estimated the posterior distributions of the time constant, plateau parameter, and intercept, which reflect our updated understanding of these parameters after considering the data. The two independent models included R2* or MTsat as the dependent variables, and the plateau parameter, time constant, and intercept, as independent variables. A Gaussian likelihood function, assuming additive normally distributed measurement noise and biological variation, was used to model the relationship between age and R2* or MTsat.

For R2*, we assigned:

– A uniform prior for the plateau parameter a (ranging from 10 to 200 (1/s)),
– A uniform prior for the intercept c (ranging from 0 to 100 (1/s)),
– A Gaussian prior for the time constant b (mean of 16 years, standard deviation of 25 years), based on values reported in previous studies for the lifespan trajectory of iron^56^.

For MTsat, we used:

– A uniform prior for the plateau parameter a (ranging from 0 to 10p.u.),
– A uniform prior for the intercept c (ranging from 0 to 10p.u.),
– A Gamma distribution for the time constant b (with parameters b=2yr, p=1yr), designed to cover a range of plausible time constants between 0.25 and 10 years based on prior research on the lifespan trajectory of myelination in humans and macaques^81^.

Model Comparisons and Evidence - To evaluate how well certain values of the time constant fit the data, we computed Bayes factors. For R2*, the Bayes factor for a time constant of 27 years was approximately 1.2, indicating weak evidence for this value. For MTsat, the Bayes factor for a time constant of 1.7 years was around 2, suggesting moderate evidence for this value.

To assess differences between regions, we compared the posterior distributions of the time constants. This comparison was done using the contrast of posterior distributions as described by ^80^, allowing us to identify how the time constants varied between different brain regions.

## Acknowledgments

This project has been funded by the Max Planck Society under the inter-institutional funds of the president for the Evolution of Brain Connectivity Project. Nikolaus Weiskopf received funding from the European Research Council under the European Union’s Seventh Framework Programme (FP7/2007-2013) / ERC grant agreement number 616905, and the European Union’s Horizon 2020 research and innovation programme under the grant agreement number 681094. This project received funding from the Deutsche Forschungsgemeinschaft (DFG, German Research Foundation)—project no. 347592254 (WE 5046/4-2 and KI 1337/2-2).

We are grateful to Caroline Jantzen, Felix Büttner, Niklas Alsleben and Franziska Zahn for their help with sample preparation and scanning and segmentation, and to Lenka Vaculčiaková for help with setting up the scan protocol. We thank the Ministère de l’Enseignement Supérieur et de la Recherche Scientifique and the Office Ivoirien des Parcs et Réserves for permitting the study in Côte d’Ivoire, Uganda Wildlife Authority and Ugandan National Council for Science and Technology for permitting the study in Uganda. Thanks to the staff of the Tai Chimpanzee Project and Budongo Conservation Field Station, Kolmarden Zoo, Chester Zoo and Twycross Zoo for their commitment.

We thank Robert Turner for feedback on previous versions of the manuscript.

## Author contributions

Conceptualization: MC, EK, IL, SE, KK, NW;

Methodology: MC, EK, IL, KP, LE, TG, CC, RW, SH, FB, NW, CJ, DR, PM, DC, RM

Software: MC, LE, NW, IL, SH;

Formal analysis: MC, IL, PM, DR, SH, LE;

Investigation: MC, EK, IL, FB, CJ, SH, PM, DR, CC, RW, NW;

Resources: NW, RW, CC, EBC Consortium;

Data Curation: MC, IL, PM, DR, SH; Writing - Original Draft: MC;

Writing - Review & Editing: MC, EK, IL, FB, CJ, KP, LE, SE, KK, SH, PM, DR, TG, RM, DC, CC, RW, NW;

Visualization: MC; Supervision: EK, NW, SH;

Project administration: NW, CC;

Funding acquisition: NW, CC, RW

## Data and materials availability

All data, code, and materials used in the analyses must be available in some form to any researcher for purposes of reproducing or extending the analyses. Include a note explaining any restrictions on materials, such as materials transfer agreements (MTAs). Include accession numbers to any data relevant to the paper and deposited in a public database; include a brief description of the dataset or model with the number. The DMA statement should include the following: “All data are available in the main text or the supplementary materials.”

## EBC Consortium members

Bala Amarasekaran^1^, Alfred Anwander^2^, Caroline Asiimwe^3^, Penelope Carlier^4^, Maelig Chauvel^5^, Julian Chantrey^6^, Catherine Crockford^7, 8, 4^, Tobias Deschner^9, 10^, Ariane Düx^11, 12^, Luke J. Edwards^5^, Cornelius Eichner^2^, Pawel Fedurek^13, 3^, Angela D. Friederici^2^, Zoro B. GoneBi^14, 4^, Tobias Gräßle^12, 11^, Philipp Gunz^15^, Jennifer E. Jaffe^11, 4^, Carsten Jäger^5^, Anna Jauch^5^, Evgeniya Kirilina^5^, Fabian H. Leendertz^16, 11^, Ilona Lipp^5^, Matyas Liptovs---zky^17^, Patrice Makouloutou Nzassi^18^, Matthew McLennan^19^, Sophie Moittie^17^, Torsten Møller^20^, Markus Morawski^21^, Karin Olofsson-Sannö^22^, Kerrin Pine^5^, Andrea Pizarro^1^, Kamilla Pleh^11, 4^, Jessica Rendel^17^, Liran Samuni^23, 4^, Lara Southern^9, 10^, Mark Stidworthy^24^, Tanguy Tanga^18, 10^, Steve Unwin^25^, Sue Walker^26^, Nikolaus Weiskopf^5^, Roman M. Wittig^7, 8, 4^, Kim Wood^27^, Klaus Zuberbuehler^28, 3^.

^1^Tacugama Chimpanzee Sanctuary, Freetown, Sierra Leone, ^2^Department of Neuropsychology, Max Planck Institute for Human Cognitive and Brain Sciences, Leipzig, Germany, ^3^Budongo Conservation Field Station, Masindi, Uganda, ^4^Tai Chimpanzee Project, CSRS, Abidjan, Cote d’Ivoire, ^5^Department of Neurophysics, Max Planck Institute for Human Cognitive and Brain Sciences, Leipzig, Germany, ^6^Veterinary Pathology and Preclinical Sciences, University of Liverpool, UK, ^7^Department of Human Behavior, Ecology and Culture, Max Planck Institute for Evolutionary Anthropology, Leipzig, Germany, ^8^Ape Social Mind Lab, Institute of Cognitive Science Marc Jeannerod, UMR 5229, CNRS, Lyon, France, ^9^Institute for Cognitive Sciences, University of Osnabrueck, Germany, ^10^Ozouga, Loango Chimpanzee Project, Loango National Park, Gabon, ^11^Robert Koch Institute, Berlin, Germany, ^12^Helmholtz Centre for Infection Research, Greifswald, Germany, ^13^School of Psychology, University of Stirling, UK, ^14^Department of Bioscience, University Felix Houphouet-Boigny, Abidjan, Cote d’Ivoire, ^15^Department of Human Origins, Max Planck Institute for Evolutionary Anthropology, Leipzig, Germany, ^16^Helmholtz Institute for One Health, Greifswald, Germany, ^17^Twycross Zoo, UK, ^18^Institut de Recherche en Ecologie Tropicale, Libreville, Gabon, ^19^Bulindi Chimpanzee and Community Project, School of Social Sciences, Oxford Brookes University, UK, 20Kolmarden Zoo, Sweden, ^21^Leipzig University, Paul Flechsig Institute of Brain Research, Leipzig, Germany, 22National Veterinary Institute, Uppsala, Sweden, 23Human Evolutionary Biology, Harvard University, Cambridge, USA, 24International Zoo Veterinary Group, Keighley, UK, 25Wildlife Health Australia, Sydney, Australia, 26Chester Zoo, UK, 27Welsh Mountain Zoo, UK, 28Institute of Biology, University of Neuchatel, Switzerland

## Supplementary Materials

### Supplementary Text

#### Lifespan trajectories of myelination and iron content

To model the age-dependent variation of iron and myelin content in chimpanzee cortical regions (foot, hand, and face), we employed an exponential saturation model to capture the dynamics of R2* (reflecting iron and myelin content) and MTsat (reflecting myelination;):

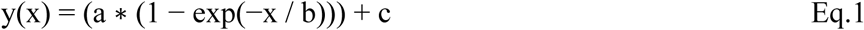

where y(x) represents the level of iron or myelin estimated from qMRI parameters at a certain age; x represents the age; a represents the total increase of selected metric (R2* or MTsat) across the entire lifespan, giving together with c the maximal value reached in old age; b represents the characteristic time constant, such that the saturation rate of lifespan changes is 1/b; and c represents the value at birth. This model was previously employed by ^56^ to model the lifespan trajectory of iron accumulation in the human brain.

Given the relatively small sample size and sparse sampling across the chimpanzee lifespan, we applied a Bayesian approach to estimate the posterior distributions of the model parameters, which incorporates prior knowledge and provides a full posterior distribution for each fitted parameter allowing for more robust parameter estimation and uncertainty quantification than conventional least-squares regression.

Using a Gaussian likelihood function, assuming additive normally distributed measurement noise, we estimated the time constants for these processes. Detailed model specifications and implementation in PyMC are provided in the methods section.

As can be seen in Supplementary Material-2, the 68% credible interval (the range within which the parameter value lies with 68% probability, given the data and the model) for the time constant b of R2* was estimated to be between 14 and 45 years, with a maximum probability at 27.5 years, which is larger than those typically reported in human cortical regions (20 years for the motor cortex) ^56^ha. For MTsat, the 68% credible interval for the time constant b ranged between 1 and 3 years, and no significant differences were observed across individual cortical regions. Posterior distributions of time constants for both R2* and MTsat across regions showed similar patterns, with contrasts symmetrically centered around zero (see Supplementary materials-2, 1.), suggesting comparable lifespan trajectories in the foot, hand, and face regions.

##### S1. Histological staining of one brain slice of a chimpanzee individual to determine the presence of Betz cells in the precentral gyrus

**Fig. S1.**
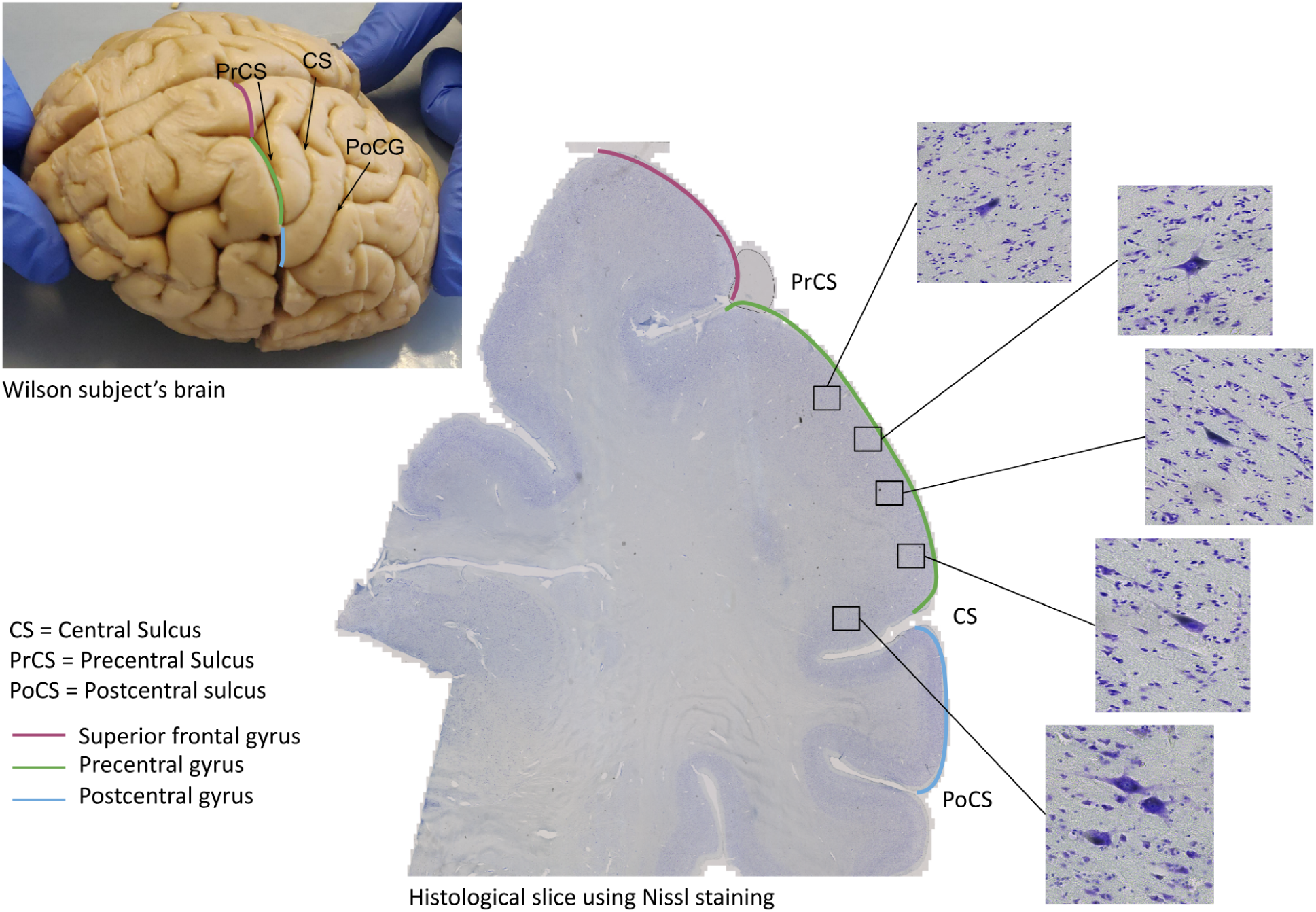
Histological staining of one brain slice of a chimpanzee individual to determine the presence of Betz cells in the precentral gyrus. Histological staining of one brain slice of a chimpanzee individual to determine the presence of Betz cells in the precentral gyrus. To investigate cortical cytoarchitecture, Nissl staining was applied. Sections were mounted on glass slides coated with Poly-L-Lysin, air dried, rehydrated and fixed in 96% ethanol for 30 min, stained with 0.1% acetate buffered cresyl violet, washed in distilled water and differentiated in graded ethanol steps. A range of Betz cells – markers of the primary motor cortex (Brodmann area 4) – could be observed along the precentral gyrus (green line).

##### S2. Lifespan trajectories of the mean R2* (left) and MTsat (right) values in the foot, hand, and face motor cortical fields in chimpanzees

**Fig. S2.**
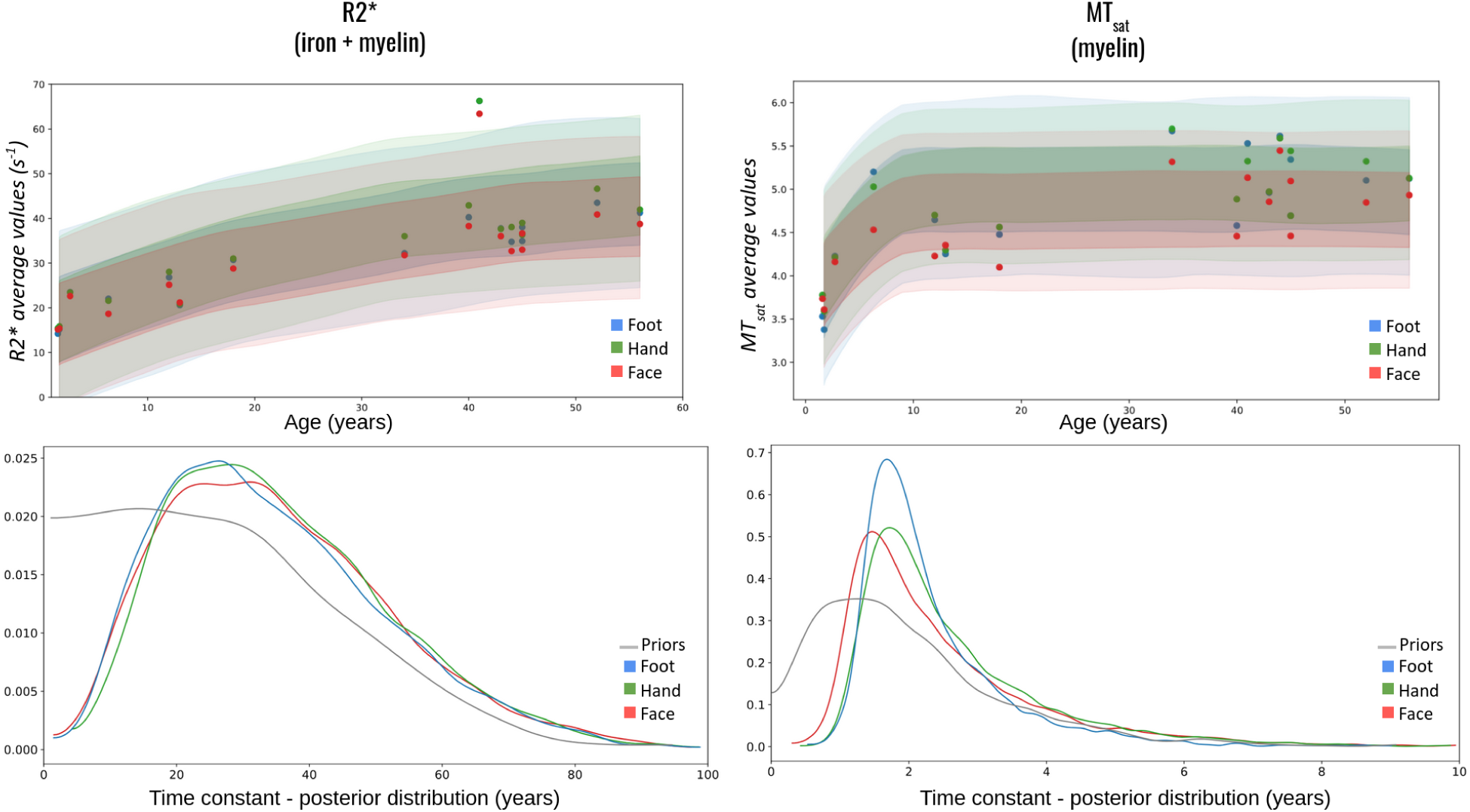
Lifespan trajectories of the mean R2* (left) and MTsat (right) values in the foot, hand, and face motor cortical fields in chimpanzees. Lifespan trajectories of the mean R2* (left) and MTsat (right) values in the foot, hand, and face motor cortical fields in chimpanzees. Top row: Applying the Hallgren and Sourander exponential saturation model ^56^ to the data for foot (blue), hand (green) and face (red) cortical fields. Shaded areas indicate the 68%- and 95%-highest density intervals. Bottom row: Identification of the time constant variation between the different cortical fields, this comparison was done using the contrast of posterior distributions as described by ^80^. Note the truncated y-axis in the top right figure.

##### S3 - Additional Analyses using independent functional validation of motor cortical field delineation

To further validate the anatomical delineation of motor cortical fields used in the main analyses, we performed an additional analysis using independent functional motor activation maps from the Human Connectome Project (HCP).

Group-level fMRI motor task activation maps were obtained from the HCP dataset (first 900 subjects from the second public release) using the Connectome Workbench platform (Marcus et al., 2011). These maps include contrasts corresponding to movements of the hand, foot, and tongue. The group-averaged activation maps were projected from the HCP surface space to the fsaverage surface space and converted into cortical labels representing the three motor representations.

Because the HCP motor task involves finger tapping, which is known to engage both the primary motor cortex (M1) and the primary somatosensory cortex (S1), the resulting activation maps extend across both the precentral and postcentral gyri. To ensure consistency with the analyses presented in the main manuscript, which specifically target motor cortex microstructure, the functional ROIs were restricted to the precentral gyrus (Brodmann area 4, BA4). The boundaries of BA4 were defined using established anatomical landmarks. The posterior boundary (BA4/BA3) was defined at the fundus of the central sulcus, corresponding to the transition between the precentral (motor) and postcentral (somatosensory) gyri. The anterior boundary (BA4/BA6) was defined at the crest of the precentral sulcus, separating primary motor cortex from premotor cortex. These criteria are supported by converging cytoarchitectonic and myeloarchitectonic studies in humans and non-human primates and are consistent with the approach used in the main manuscript. In addition, these boundaries were verified in one post-mortem brain using histological criteria, including differences in cortical lamination and the presence of Betz cells in BA4.

The functional ROIs were applied to the cortical surfaces of the human participants included in this study (7T cohort, n = 10; 3T cohort, n = 44). For each subject, R2* and MTsat values were sampled across cortical depth between the pial surface and the white matter boundary using the same procedure as described in the main manuscript. Microstructural values were extracted within each functional ROI (hand, foot, and face/tongue representations), and analyses were performed following the same statistical procedures as in the main study.

###### Interpretation

The comparisons of microstructural values across motor representations obtained using functionally defined ROIs are shown in Supplementary Figure S4. Both R2* and MTsat values differed across motor cortical fields in a manner consistent with the main results, with the hand representation exhibiting higher values relative to the face/tongue and foot representations in both human cohorts.

Hemispheric differences within the hand motor field are shown in Supplementary Figure S5. When using functionally defined ROIs, hemispheric asymmetries were consistent with those reported in the main manuscript, indicating that the inclusion of somatosensory cortex in broader functional ROIs can influence the detection of localized hemispheric effects.

**Figure S3.**
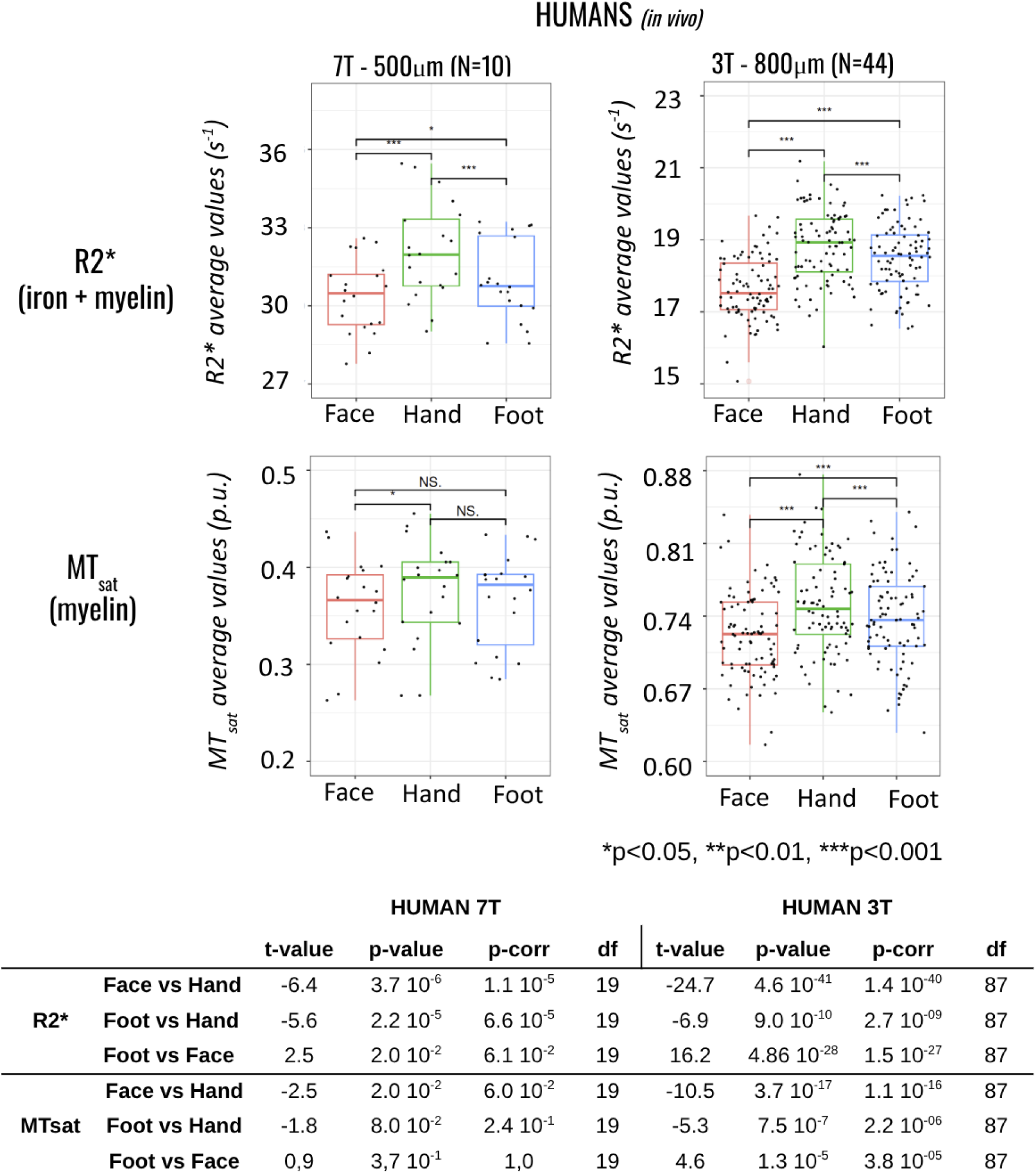
Microstructural differences across motor representations using functional ROIs. R2* and MTsat values averaged across functionally defined motor cortical fields in the human datasets. (top) Boxplots show R2* values (top row) and MTsat values (bottom row) extracted from face/tongue (red), hand (green), and foot (blue) motor cortical regions defined using Human Connectome Project functional activation maps. Results are presented for two independent human cohorts acquired at different field strengths: 7T (n = 10; left column) and 3T (n = 44; right column). (bottom) Statistical test results comparing qMRI metrics between the different cortical fields in humans. Corrections for multiple comparisons were made using the Bonferroni correction (p-corr). Differences between motor cortical fields were assessed using paired t-tests. Boxes represent the 25th and 75th percentiles; horizontal lines indicate the median; whiskers represent the most extreme data values. Note the truncated y-axes. *p < 0.05, **p < 0.01, ***p < 0.001.

**Figure S4.**
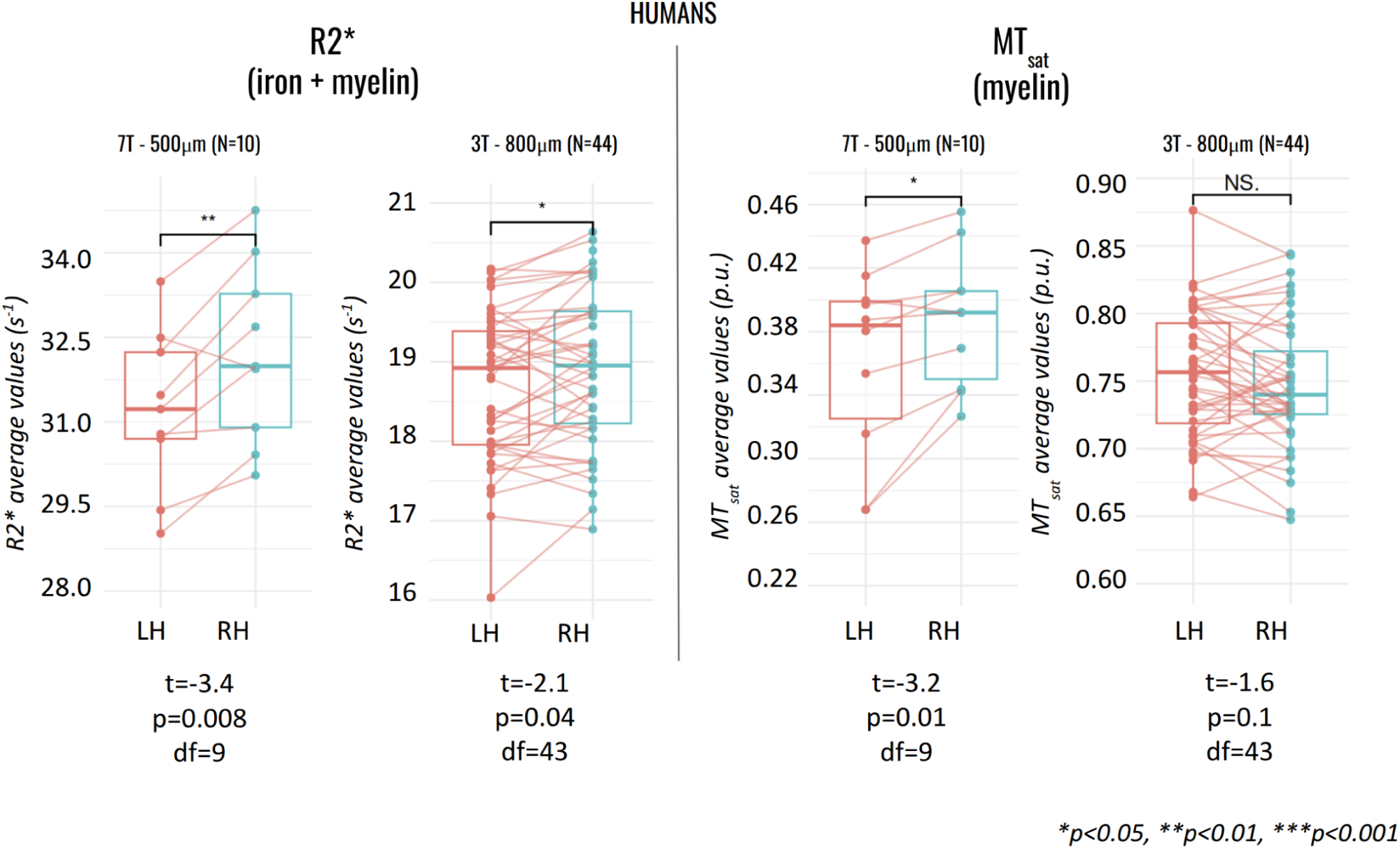
Hemispheric asymmetry in the functionally defined hand motor field. R2* and MTsat values averaged within the functionally defined hand motor cortical field for the left and right hemispheres in the human datasets. Boxplots show R2* values (left) and MTsat values (right) for participants scanned at 7T (n = 10) and 3T (n = 44). The hand region was defined using Human Connectome Project motor task activation maps. Boxes represent the 25th and 75th percentiles; horizontal lines indicate the median; whiskers represent the most extreme values excluding outliers. Individual data points are shown and paired measurements from the same participants are connected by lines. Differences between hemispheres were assessed using paired t-tests. Note the truncated y-axes. *p < 0.05, **p < 0.01, ***p < 0.001.

These analyses provide an independent functional validation of the anatomical delineation used in the main study. The results confirm that the observed microstructural differences across motor cortical representations, as well as hemispheric asymmetries within the hand motor field, are robust when motor regions are defined using independent functional activation maps constrained to primary motor cortex.

##### S4 - Effect of age distribution in the chimpanzee cohort

To address the differences in age distribution between chimpanzee and human cohorts, we performed an additional analysis restricting the chimpanzee sample to individuals above early developmental stages. Specifically, we excluded chimpanzees younger than 6 years of age (namely Lukule, Rasta and Flag), as this period corresponds to ongoing brain maturation, including myelination processes in motor cortical regions.

All analyses were then repeated on this restricted cohort, including (i) comparisons of microstructural values across motor cortical fields (see figure S5) and (see figure S6) hemispheric asymmetry analyses within the hand motor representation.

**Figure S5.**
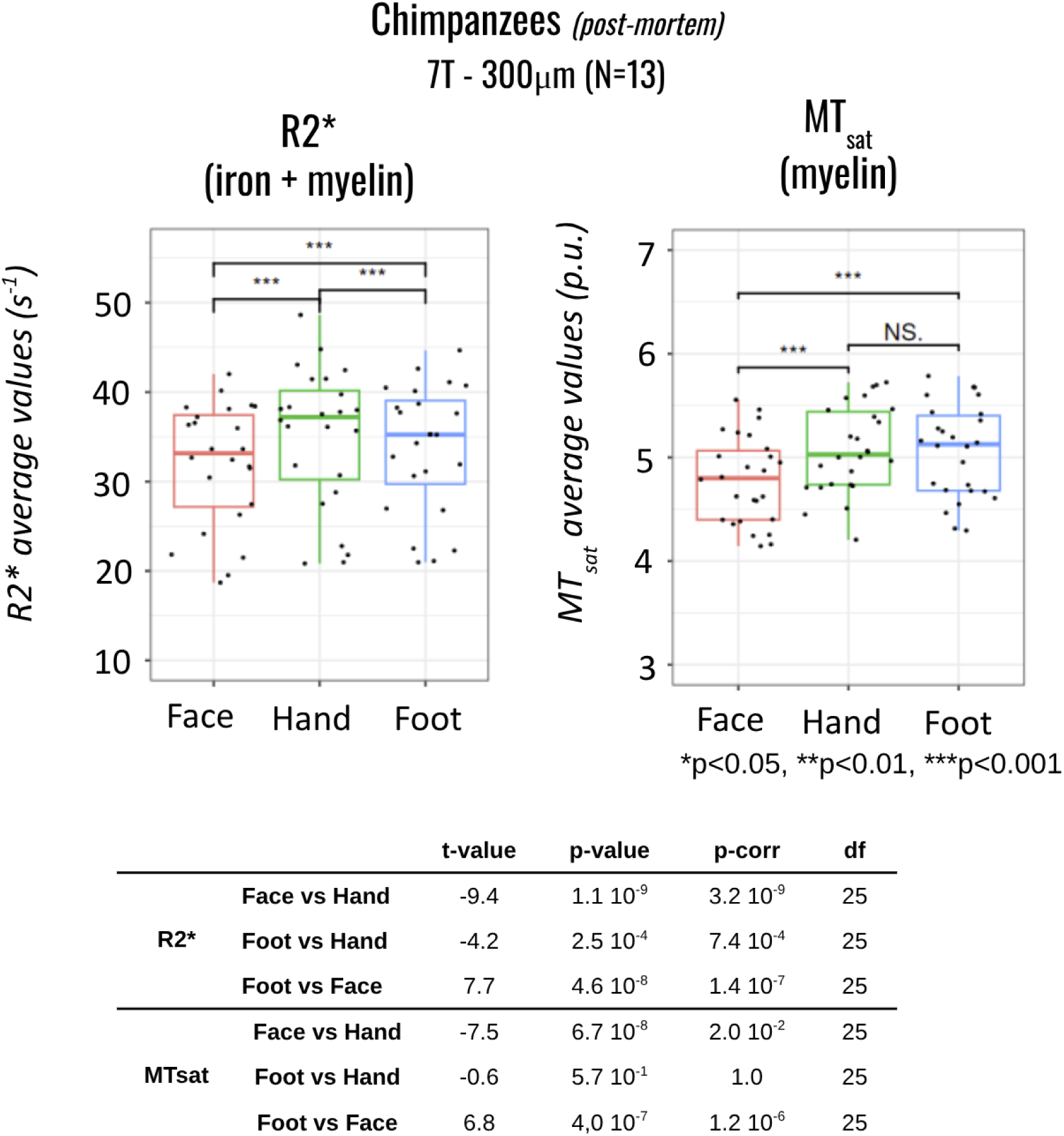
Microstructural differences across motor representations in adult chimps (over 6 years old). R2* and MTsat values averaged across functionally defined motor cortical fields in the chimpanzee datasets containing only adults (n= 13, over 6 years old). (top) Boxplots show R2* values (left) and MTsat values (right) extracted from face/tongue (red), hand (green), and foot (blue) motor cortical regions. Differences between motor cortical fields were assessed using paired t-tests. Boxes represent the 25th and 75th percentiles; horizontal lines indicate the median; whiskers represent the most extreme data values. Note the truncated y-axes. *p < 0.05, **p < 0.01, ***p < 0.001. (bottom) Statistical test results comparing qMRI metrics between the different cortical fields in adult chimpanzees. Corrections for multiple comparisons were made using the Bonferroni correction (p-corr).

**Figure S6.**
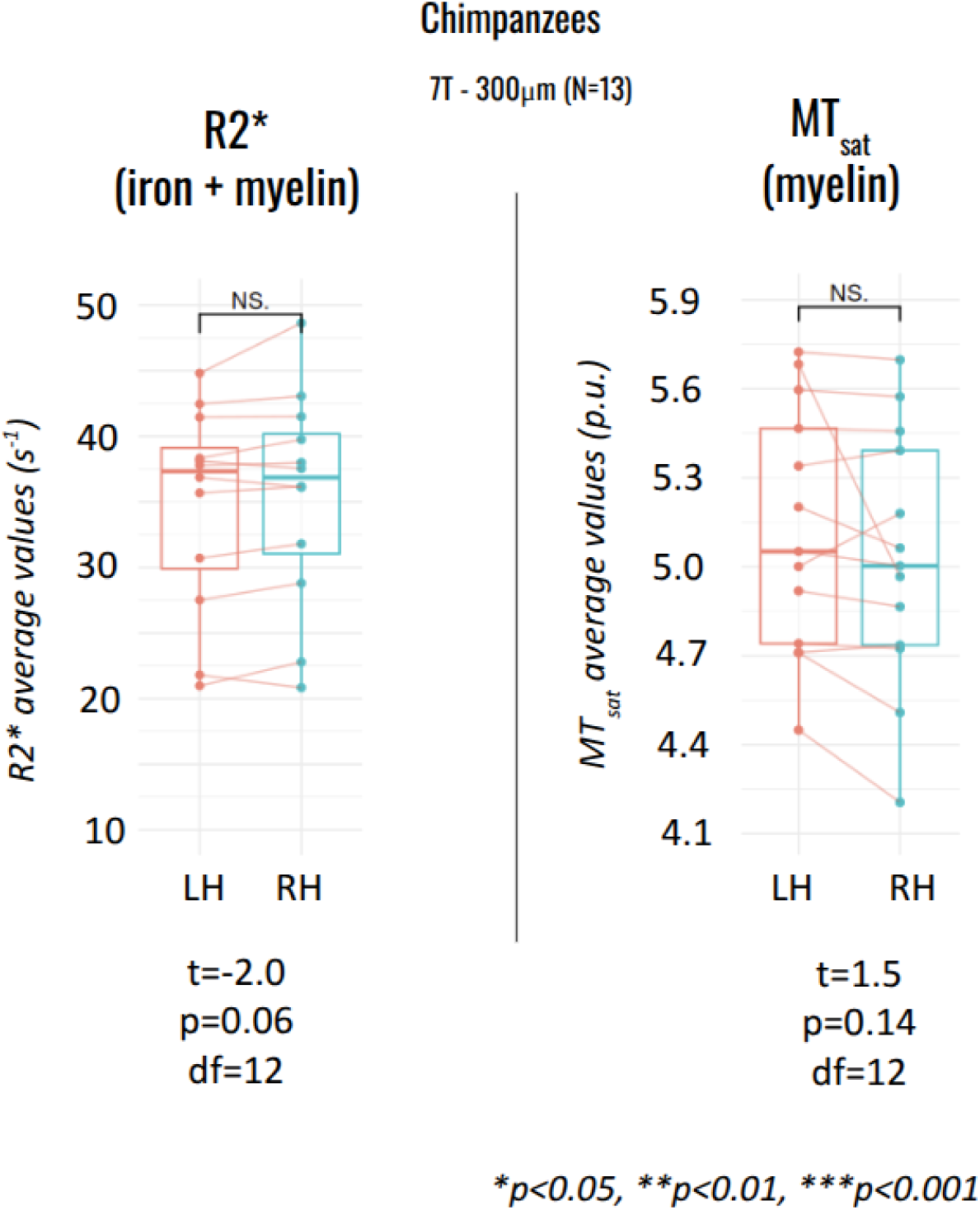
Hemispheric asymmetry in the hand motor field in adult chimpanzees. R2* and MTsat values averaged within the defined hand motor cortical field for the left and right hemispheres in the chimpanzee dataset restricted to adult chimps (N=13, over 6 years old). Boxplots show R2* values (left) and MTsat values (right) for chimpanzees scanned at 7T. Boxes represent the 25th and 75th percentiles; horizontal lines indicate the median; whiskers represent the most extreme values excluding outliers. Individual data points are shown and paired measurements from the same participants are connected by lines. Differences between hemispheres were assessed using paired t-tests. Note the truncated y-axes. *p < 0.05, **p < 0.01, ***p < 0.001.

